# Dopaminoceptive D1 and D2 neurons in ventral hippocampus arbitrate approach and avoidance in anxiety

**DOI:** 10.1101/2023.07.25.550554

**Authors:** Arthur Godino, Marine Salery, Angelica M. Minier-Toribio, Vishwendra Patel, John F. Fullard, Eric M. Parise, Freddyson J. Martinez-Rivera, Carole Morel, Panos Roussos, Robert D. Blitzer, Eric J. Nestler

**Affiliations:** Nash Family Department of Neuroscience & Friedman Brain Institute, Icahn School of Medicine at Mount Sinai, New York, NY 10029, USA; Department of Psychiatry & Friedman Brain Institute, Icahn School of Medicine at Mount Sinai, New York, NY 10029, USA; Department of Pharmacological Sciences, Icahn School of Medicine at Mount Sinai, New York, NY 10029, USA; Department of Genetics and Genomic Sciences & Icahn Genomics Institute, Icahn School of Medicine at Mount Sinai, New York, NY 10029, USA; Center for Disease Neurogenomics, Icahn School of Medicine at Mount Sinai, New York, NY 10029, USA; Mental Illness Research, Education and Clinical Centers, James J. Peters VA Medical Center, Bronx, NY 10468, USA

## Abstract

The hippocampus ^1–7^, as well as dopamine circuits ^8–11^, coordinate decision-making in anxiety-eliciting situations. Yet, little is known about how dopamine modulates hippocampal representations of emotionally-salient stimuli to inform appropriate resolution of approach *versus* avoidance conflicts. We here study dopaminoceptive neurons in mouse ventral hippocampus (vHipp), molecularly distinguished by their expression of dopamine D1 or D2 receptors. We show that these neurons are transcriptionally distinct and topographically organized across vHipp subfields and cell types. In the ventral subiculum where they are enriched, both D1 and D2 neurons are recruited during anxiogenic exploration, yet with distinct profiles related to investigation and behavioral selection. In turn, they mediate opposite approach/avoidance responses, and are differentially modulated by dopaminergic transmission in that region. Together, these results suggest that vHipp dopamine dynamics gate exploratory behaviors under contextual uncertainty, implicating dopaminoception in the complex computation engaged in vHipp to govern emotional states.

## Main

Anxiety promotes adaptive safety and survival reactions, but when inappropriate to the level of threat contributes to psychiatric disorders ^12^. Both healthy and dysregulated emotional processing related to unconditioned fear expression and approach/avoidance conflict resolution have long implicated the hippocampus ^1,2^, and especially its ventral pole (vHipp) in rodents or anterior hippocampus in humans ^3–5^. Current theoretical models postulate that vHipp computes diverse – contextual and internal – input signals and enables appropriate behavioral selection by driving relevant, mostly parallel ^13,14^ output projections, in turn flexibly arbitrating approach or avoidance ^6,7^. Yet, the exact combinatorial logic at play remains unclear: while sparse and heterogeneous responses to anxiogenic stimuli have been observed in vHipp ^15–17^, phenotypical markers favoring the recruitment of specific neurons or subsets of neurons into those representations are still lacking.

Dopamine axons from the midbrain innervate vHipp, comparatively more densely in the ventral CA1 (vCA1) and adjacent subiculum (vSub) ^18–20^, where most hippocampal projection neurons reside ^21^. vHipp also exhibits topographically organized expression of G-protein-coupled dopamine receptors ^22–24^, with *Drd1*/D1 and *Drd2*/D2 receptors seemingly separated between largely non-overlapping populations ^25^. This is akin to medium spiny neurons (MSNs) of striatum, whose segregated D1 or D2 expression is integral to proper dopamine signal processing and behavioral execution ^26–29^. In striatum and other brain regions, dopamine signals carry critical information for motivated approach behaviors and reward learning on one hand ^30^, and avoidance and aversive learning on the other ^10,31^. Consequently, dysfunction in dopamine pathways is associated with anxiety traits and disorders ^8,9,11,32^ and, although those behavioral functions are likewise intimately linked to vHipp activity ^6,7^, specific insight into dopaminergic modulation of vHipp networks has remained surprisingly elusive.

Therefore, we here investigate the influence of dopaminoceptive signaling within vHipp on anxiety-related approach and avoidance behaviors. We hypothesize that the D1/D2 status of singular vHipp neurons might represent an important determinant in the underlying circuit-level computation – a purposely broad initial premise given the near non-existent literature on vHipp dopaminoception.

### D1- and D2-expressing cells are topographically organized in vHipp

To first visualize D1- and D2-expressing cell bodies in vHipp, we capitalized on well-validated BAC transgenic lines expressing Cre recombinase under the respective control of the *Drd1* (D1-Cre) or *Drd2* (D2-Cre) gene promoters, crossed to ^fl/fl^eGFP::L10a reporter mice ^33^. On representative vHipp sections, these two cell types exhibit clearly distinct histological organization (Fig. 1). In the dentate gyrus (DG), D1 cells are mostly located in the granule cell layer, while D2 cells are located almost exclusively in the polymorphic layer or hilus – the latter being consistent with published litterature ^25,34,35^. In CA3, which is smaller at such caudal coordinates, sparser labeling was observed for D2 cells than for D1 cells across layers. In vCA1, sparse D2 and even sparser D1 cells were found in the stratum lacunosum moleculare, especially at the radiatum-lacunosum moleculare (R-LM) border, and in the stratum oriens, across the dorso-ventral axis. In the vCA1 pyramidal cell layer, gradual enrichment of both D1 and D2 cells emerges ventral to the rhinal fissure, along with the diffuse ^21^ transition from vCA1 to vSub. Together, these semi-quantitative observations indicate a precise topographical organization of D1 and D2 cells across vHipp subfields and layers, most notably in the DG and in the caudal-most parts of vCA1/vSub.

**Fig. 1:**
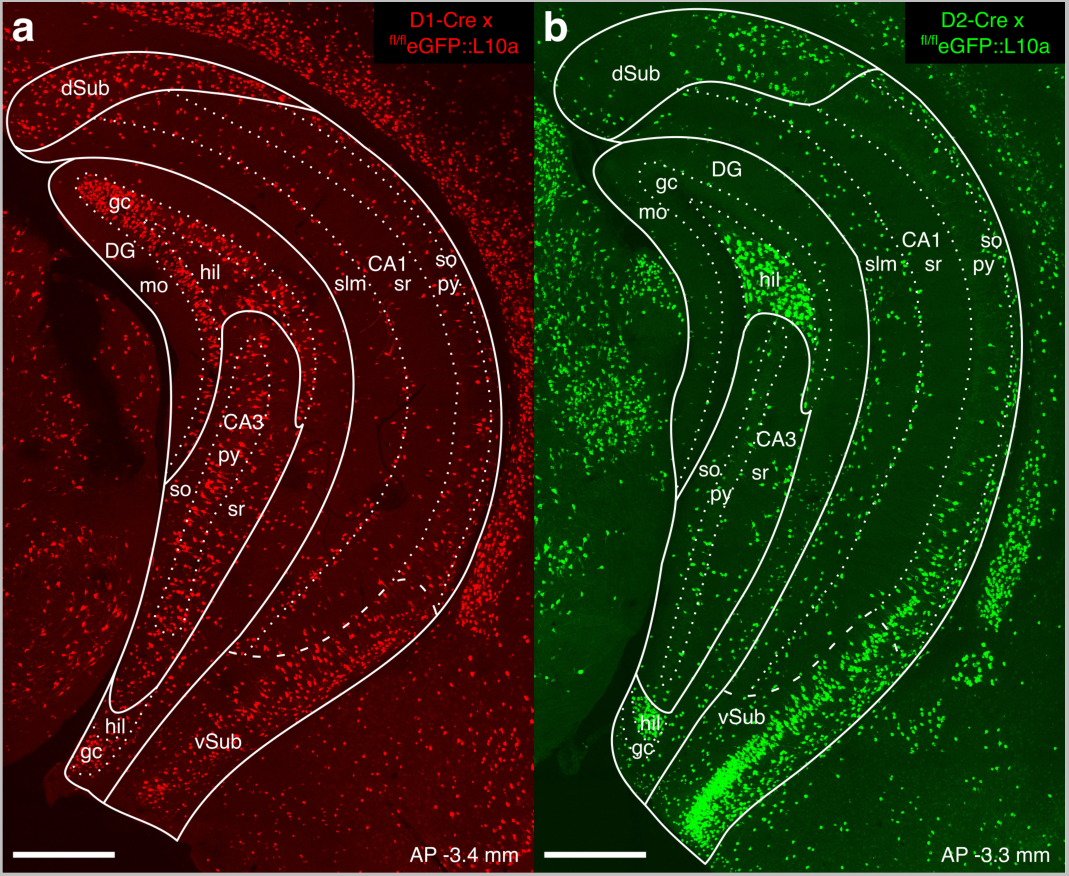
Topography of vHipp D1 and D2 cells. Representative confocal images of the vHipp of a D1-Cre (**a**) and D2-Cre (**b**) x ^fl/fl^eGFP::L10a male mouse. D1 cells are GFP-positive, shown here in pseudo-red color for clarity. Scale bars 500 µm. Abbreviations: AP antero-posterior from Bregma, DG dentate gyrus, mo molecular layer, gc granule cell layer, hil hilus, slm stratum lacunosum moleculare, sr stratum radiatum, py pyramidal cell layer, so stratum oriens.

### vHipp D1 and D2 cells are segregated across and within neuronal cell types

We next more thoroughly phenotyped vHipp dopaminoceptive cells via single-nuclei RNA-sequencing (snRNAseq) of D1- and D2-expressing nuclei (Fig. 2a) isolated by fluorescence-activated nuclei-sorting (FANS, Extended Data Fig. 1), each of which represented ~6% of the total nuclei counts (Extended Data Fig. 1c). From a merged dataset containing transcriptomes from all captured D1 and D2 nuclei, unsupervised dimensionality reduction approaches identified 21 clusters (Fig. 2b and Extended Data Fig. 2a,b), which were further annotated by comparison (Extended Data Fig. 2d and Supplementary Information Table 1) to publicly available single-cell RNAseq databases ^36–40^. With the exception of a very sparse (0.4%) cluster of astrocytes, vHipp D1 and D2 cells were all neuronal. GABAergic interneurons were over-represented (53.3% of total) compared to whole hippocampus snRNAseq datasets where they only account for <10% of all cells ^41^. Yet, glutamatergic neurons represented 45.1% of all D1 or D2 cells in vHipp, in stark contrast to more dorsal parts of hippocampus where – barring D2-positive hilar mossy cells – D1 or D2 cells are almost exclusively interneurons ^34,35,42^. While GABAergic clusters readily mapped to canonical neuropeptide-defined interneuron cell types ^40^, glutamatergic pyramidal neuron classification was not as clear-cut: we hypothesize that pyramidal neuron clusters might generally correspond to projection-specific vCA1/vSub populations.

**Fig. 2:**
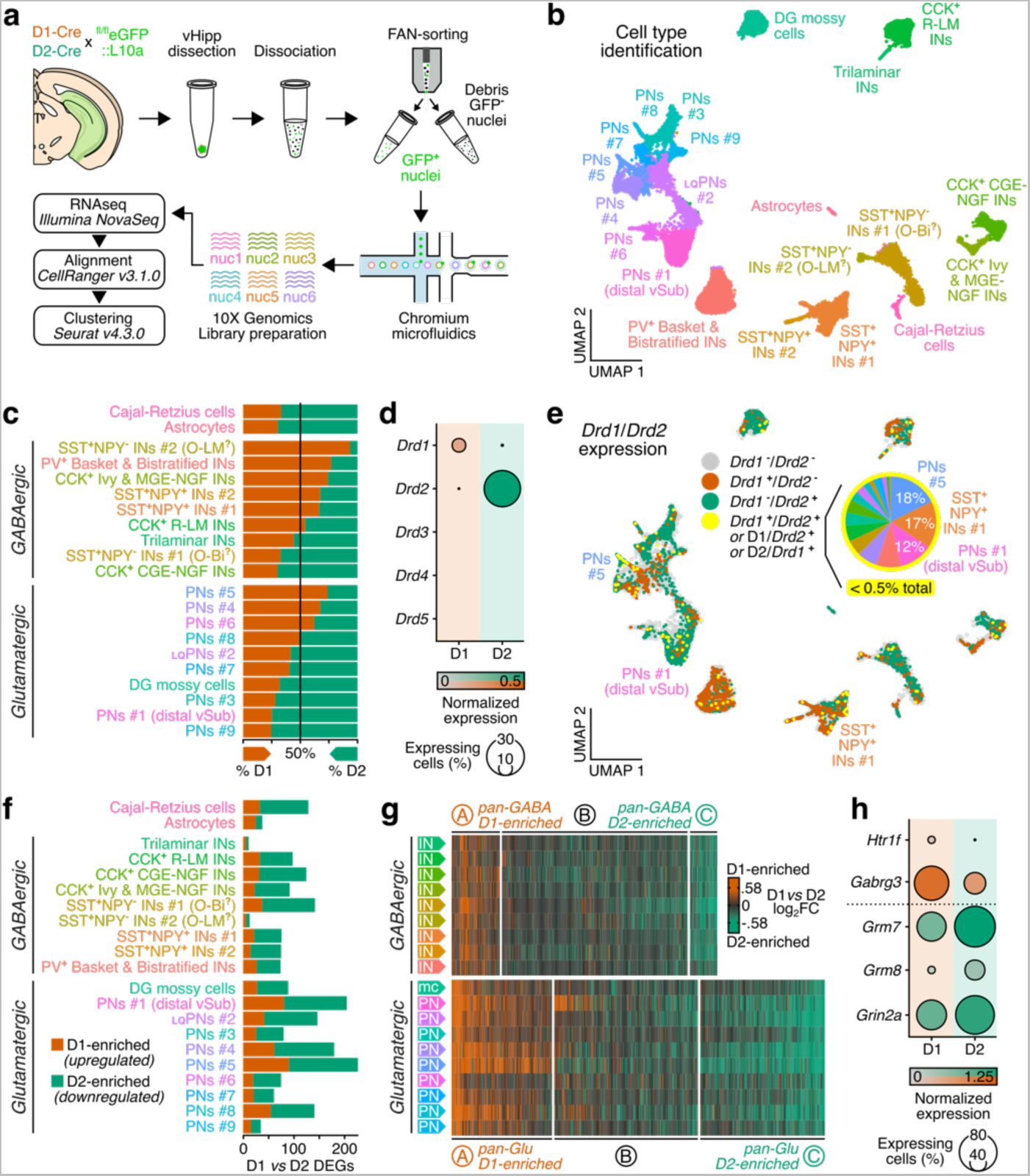
Transcriptional phenotypes of vHipp D1 and D2 cells. **a**, Workflow for snRNAseq of vHipp from male D1-Cre and D2-Cre x ^fl/fl^eGFP::L10a (n = 4/genotype). **b**, Clustering and UMAP reduction across all samples (n = 11,452 D1 and n = 18,158 D2 nuclei) followed by cluster-cell type annotation. **c**, Proportion of nuclei originating from FANS-isolated D1 and D2 samples per cluster, normalized to total D1 and D2 nuclei counts. **d**, Expression of *Drd1*, *Drd2*, *Drd3*, *Drd4* and *Drd5* dopamine receptor genes across D1 and D2 nuclei. **e**, Expression of dopamine receptor *Drd1* and *Drd2* genes in individual nuclei. Nuclei are considered as co-expressing D1 and D2 receptors (yellow) either if both *Drd1* and *Drd2* are detected, if *Drd1* is detected in a D2-sorted nucleus or if *Drd2* is detected in a D1-sorted nucleus. Insert shows the repartition of the small (<0.5%) population of D1-D2 co-expressing nuclei across clusters. **f**, Number of differentially expressed genes (DEGs, >20% expression change and FDR-adjusted *p* < 0.05) between D1 and D2 nuclei per individual cell type clusters. **g**, Union heatmaps and hierarchical clustering of D1 *versus* D2 DEGs in different vHipp cell types. DEG clusters/patterns A and C contain genes respectively enriched in D1 and D2 nuclei across cell types (pan-GABAergic or pan-glutamatergic nuclei). **h**, Expression across D1 and D2 nuclei of neurotransmitter and neuromodulator receptor genes selected from either D1-enriched (*Htr1f*, *Gabrg3*) or D2-enriched (*Grm7*, *Grm8*, *Grin2a*) pan-GABAergic and pan-glutamatergic DEG patterns. Abbreviations: INs interneurons, PNs pyramidal neurons, _LQ_PNs low sequencing quality pyramidal neurons, CCK cholecystokinin, NPY neuropeptide Y, PV parvalbumin, SST somatostatin, O-Bi oriens – bistratified, O-LM stratum oriens / stratum lacunosum moleculare, R-LM stratum radiatum / stratum radiatum border, DG dentate gyrus, MGE medial ganglionic eminence-derived, CGE central ganglionic eminence-derived, NGF neurogliaform.

At the population level, vHipp D1 and D2 cells were segregated across individual clusters (Fig. 2c and Extended Data Fig. 2c), with some cell types comprising predominantly D1 neurons (*e.g.,* basket and bistratified interneurons) or D2 neurons (*e.g.,* DG hilar mossy cells, consistent with Fig. 1 histology). Among these clusters, very few (<0.5%) D1 and D2 vHipp cells co-expressed both dopamine receptors (Fig. 2d,e). Other dopamine receptors (*Drd3*, *Drd4*, *Drd5*) were not expressed in D1 and D2 vHipp neurons (Fig. 1d), suggesting that D3-, D4- and D5-expressing vHipp cells ^22^ might constitute yet other separate populations. Furthermore, cluster-specific differential expression analysis revealed profound transcriptional differences between D1 and D2 vHipp cells (Fig. 2f and Supplementary Information Table 2). These D1 *versus* D2 differences, however, were conserved across cell type clusters: genes enriched in one D1 pyramidal neuron or interneuron cluster were similarly enriched in all pyramidal neuron or interneuron clusters, respectively, and *vice versa* for D2-enriched genes, as evidenced by strikingly similar gene expression heatmaps and confirmed by unsupervised classification into gene expression patterns (Fig. 2g and Supplementary Information Table 3). This indicates that D1 and D2 neurons, although encompassing diverse hippocampal neuronal phenotypes, do share other common transcriptional features, including many related to synaptic transmission (Supplementary Information Table 4). Such pan-cell-types (in terms of GABAergic or glutamatergic clusters) yet D1- or D2-specific transcriptional profiles might suggest coordinated ensemble responses, for instance to G-protein-mediated neuromodulation: both principal neurons and interneurons expressing D1 receptors express higher levels of the G-protein-coupled serotonin receptor *Htr1f*, while D2 neurons express higher levels of the G-protein-coupled glutamate receptors *Grm7* and *Grm8* (Fig. 2h and Supplementary Information Table 3). Together with histology, this snRNAseq dataset illustrates that vHipp D1 and D2 cells represent distinct neuronal populations divided across cell types but with shared, denominating phenotypical properties, similarly to D1- and D2-MSNs. Because the main functional output from vHipp is coordinated by projections from vCA1/vSub ^6,7^, which we found rich in dopaminoceptive cells, we further focused on D1 and D2 neurons in these areas – referred to as vSub below (Extended Data Fig. 3).

### vSub D1 and D2 neurons are activated during exploration of anxiogenic environments

With the hypothesis that the D1/D2 status of vSub neurons might contribute to differences observed in behaviorally-relevant single-neuron activity in this region ^15–17^, we investigated the respective activation dynamics of vSub D1 and D2 neurons during anxiety-related testing. At first analysis, D1- or D2-specific *in vivo* GCaMP6s calcium imaging in freely-moving male D1- or D2-Cre mice in an elevated-plus maze (EPM) task (Fig. 3a) illustrated that both vSub D1 and D2 neurons were strongly, consistently and sustainably activated by exploration of the open arm, the anxiogenic compartment of the EPM (Fig. 3b-e and Extended Data Fig. 4a). A finer examination around head-dip events – when mice extend their head outside of the EPM apparatus, which can be interpreted as highly-anxiogenic sampling/investigation behavior and has been shown to elicit single-neuron activity in vCA1 ^17^ – detected strikingly contrasting activity patterns between the two: while D1 neurons were activated independent of whether mice will abort or continue exploration following head-dipping (although with lower signal amplitude in the latter), D2 neurons were on average only activated if mice continued exploration, and inhibited if mice demonstrated avoidance behavior (Fig. 3f,g and Extended Data Fig. 4b-d). Consistently, unsupervised clustering of individual head-dip time-series into 3 distinct activity patterns (A: transient activation, B: sustained activation, C: inhibition) confirmed stronger bias in behavioral outcomes (avoid/explore) based on calcium dynamics alone for D2 than for D1 cells (Fig. 3h). Finally, we examined whether vSub D1 or D2 calcium activity patterns around head-dips contain enough information to predict subsequent behavior, especially because signal changes occurred before movement initiation (Fig. 3f and Extended Data Fig. 4b,e). To test this idea, we put D1 and D2 head-dip time-series through a supervised machine learning paradigm capitalizing on support vector machine (SVM) classification models (Fig. 3i), which have successfully been implemented to decode vCA1 memory traces ^43^. We found that D2 signals around head-dips predicted avoid/explore decisions with good (75%) accuracy, significantly better than chance, whereas D1 signals performed barely (55%) above chance (Fig. 3j). Fully-trained SVM models could also classify events with undetermined manual annotation into categories with average activity resembling that of those with clear outcomes (Extended Data Fig. 4b,c,f), highlighting that signal computation takes place in vSub even when behavioral execution is not as clear-cut. These patterns illustrated that, although both D1 and D2 vSub neurons are recruited, subtle differences exist related to a mouse’s decision to engage in approach or avoidance during anxiogenic exploration, suggesting distinct behavioral roles of these two cell types. We further postulate that vSub D1 neurons might thus react to and encode anxiogenic features and stimuli (a detection role more related to pro-avoidance behaviors) while D2 neurons might more specifically motivate exploration under environmental conflict and investigation of those anxiogenic features (a pro-approach role, where D2 activation might “override” D1 anxiety-detecting signals). However, such a distinction is hard to resolve using correlative techniques only, especially in a task where both coexist spatially and temporally like EPM testing.

**Fig. 3:**
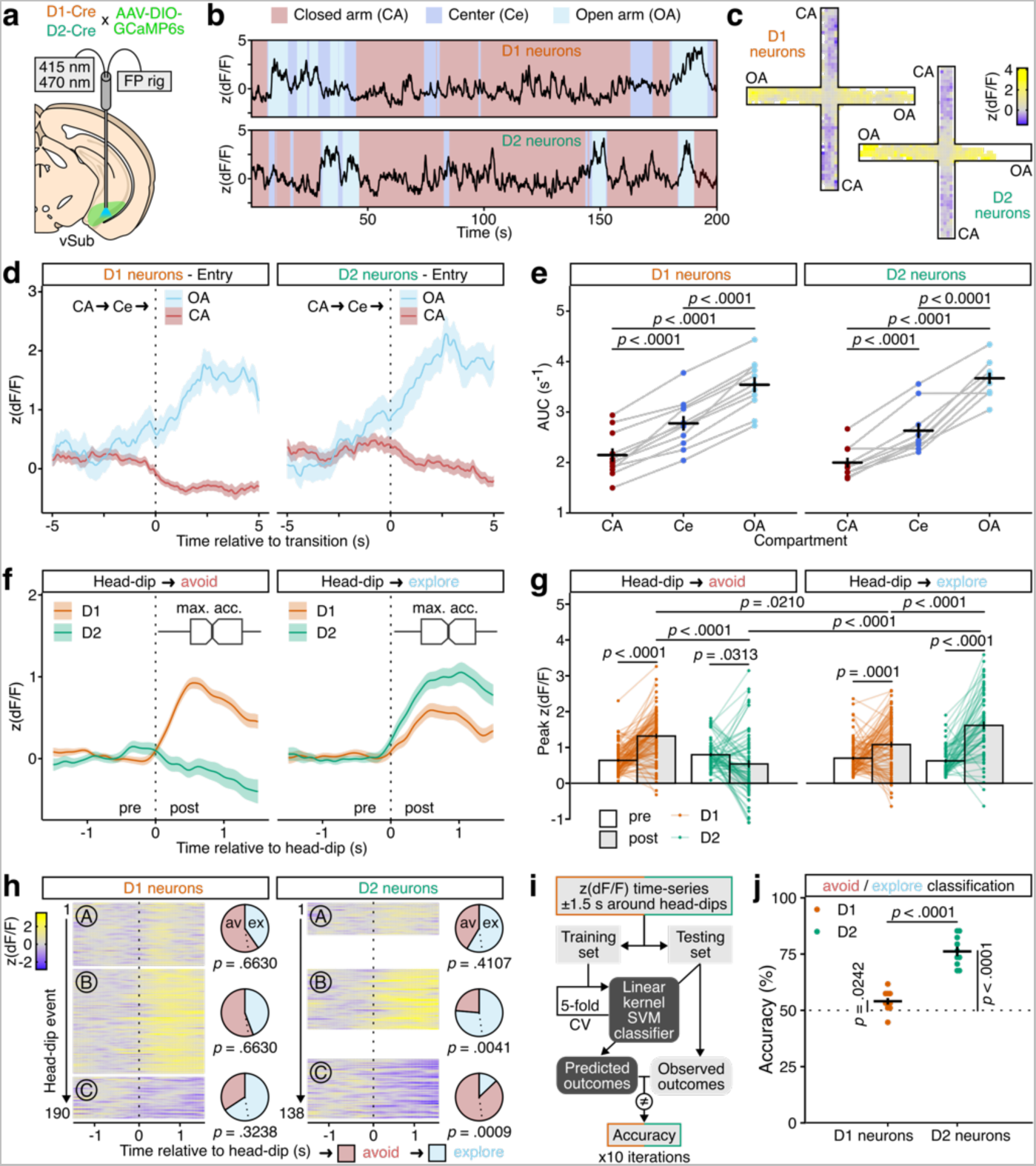
Calcium imaging of vSub D1 and D2 neuronal activity during EPM testing. **a**, Experimental schematic. Male D1-Cre (n = 11) and D2-Cre (n = 10) mice were injected with an AAV-DIO-GCaMP6s in vSub and implanted with optical fibers before recording in an EPM task. **b**, Representative GCaMP6s signal during EPM exploration from a D1-Cre (top) and D2-Cre (bottom) mouse. **c**, Spatially averaged GCaMP6s signal in the EPM for all D1-Cre (top-left) and all D2-Cre (bottom-right) mice. **d**, Average GCaMP6s signal in D1-Cre (left) or D2-Cre (right) mice during entries in the open arm (OA, blue) or closed arm as a control (CA, red). Only traces when the mouse did a complete closed arm to center (Ce) to open/closed arm are used for averaging. **e**, Average area under the curve (AUC) signal quantification by EPM compartment for D1-Cre (left) and D2-Cre (right) mice. LMM-ANOVA: compartment F_2,38_ = 255.19 *p* < 0.0001, cell type F_1,19_ = 0.10 *p* = 0.7533, compartment x cell type F_2,38_ = 2.72 *p* = 0.0785; followed by FDR-adjusted post-hoc tests. **f**, Average GCaMP6s signal in D1-Cre (red) or D2-Cre (green) mice around head-dips outside the EPM apparatus event, manually annotated based on outcome mouse behavior: aborted exploration and avoidance (left, red) or continued investigation and exploration (right, blue). Events with unclear outcome are included in analysis but shown in Extended Data Fig. 4. Boxplots represent the median time ± inter-quartile range of the times of maximal acceleration (max. acc.) after each head-dip, indicating movement initiation. **g**, Quantification of maximum (peak) GCaMP6s signal before (pre) or after (post) each head-dip event. LMM-ANOVA: cell type F_1,22.22_ = 0.27 *p* = 0.6069, outcome F_2,375.77_ = 7.06 *p* = 0.0010, pre-post F_1,385_ = 84.64 *p* < 0.0001, cell type x outcome F_2,375.77_ = 16.74 *p* < 0.0001, cell type x pre-post F_1,385_ = 0.17 *p* = 0.6775, outcome x pre-post F_2,385_ = 11.69 *p* < 0.0001, cell type x outcome x pre-post F_2,385_ = 34.37 *p* < 0.0001; followed by FDR-adjusted post-hoc tests. **h**, Unsupervised hierarchical clustering of GCaMP6s head-dip time-series for D1 (left, n = 190) and D2 (right, n = 138) neurons, pictured as signal intensity heatmaps, split into patterns A (transient activation), B (robust, sustained activation) and C (inhibition). Pie charts depict the relative proportion of avoid(av)/explore(ex) behavioral outcomes in each pattern, for each cell type, along with FDR-adjusted *p*-values corresponding to standardized Pearson’s residuals after χ^2^ tests (D1 neurons: χ^2^ = 6.68, df = 2, *p* = 0.0355; D2 neurons: χ^2^ = 45.608, df = 2, *p* < 0.0001). Dotted line indicates theoretical proportions. **i**, Conceptual schematic for supervised binary linear classification of D1 and D2 time-series using support vector machines (SVM) with 5-fold cross validation (CV). For each iteration, the whole dataset was randomly split into training (75%) and testing (25%) sets. **j**, SVM classifier accuracy for D1 and D2 time-series. Welch’s t-tests: D1 *versus* D2 t_16.16_ = −8.52 *p* < 0.0001; D1 *versus* chance (50%) t_9_ = 2.71 *p* = 0.0242; D2 *versus* chance (50%) t_9_ = 12.33 *p* < 0.0001; followed by FDR adjustment. Data represented as mean ± sem.

### vSub D1 and D2 neurons oppositely modulate exploration of anxiogenic environments

To interrogate the causal contribution of vSub D1 and D2 neurons to anxiety-related behaviors, we first used chemogenetics to express either the activating hM3Dq (Fig. 4a) or the inhibitory hM4Di (Fig. 4d) designer receptor exclusively activated by designer drugs (DREADD) selectively in either vSub D1 or D2 neurons of D1- or D2-Cre male mice, which were then artificially stimulated by systemic injection of the ligand clozapine-N-oxide (CNO) before EPM testing. This approach aimed to alter the excitability of D1 *versus* D2 cells during behavior, as photometry recordings (Fig. 3f-j) suggested that it might in fact be the relative activation balance of the two cell types that modulate behavior. CNO alone (in the absence of a DREADD) did not affect EPM behaviors (Extended Data Fig. 5). Activating D1 neurons reduced the time spent by experimental mice in the open arm (Fig. 4b), whereas activating D2 neurons increased it (Fig. 4c) together with the number of open arm entries and the average time spent in the open arm per exploration bout (Extended Data Fig. 6c), demonstrating opposite control of anxiety-related exploratory behavior. Reciprocally, inhibiting D2 neurons reduced open arm exploration time (Fig. 4f) with strong trends towards reduced open arm entries and exploration bout length (Extended Data Fig. 6f), while inhibiting D1 neurons had no significant effect (Fig. 4e and Extended Data Fig. 6e). D1 neurons might thus be sufficient but not necessary to elicit anxiety-related avoidance responses, unlike D2 neurons which appear both necessary and sufficient for counter-related approach behaviors. Testing in other anxiety-related paradigms showed consistent pro-anxiogenic effects of D1 activation in open-field exploration (Extended Data Fig. 7c) and of D2 inhibition in novelty-suppressed feeding (Extended Data Fig. 7d).

**Fig. 4:**
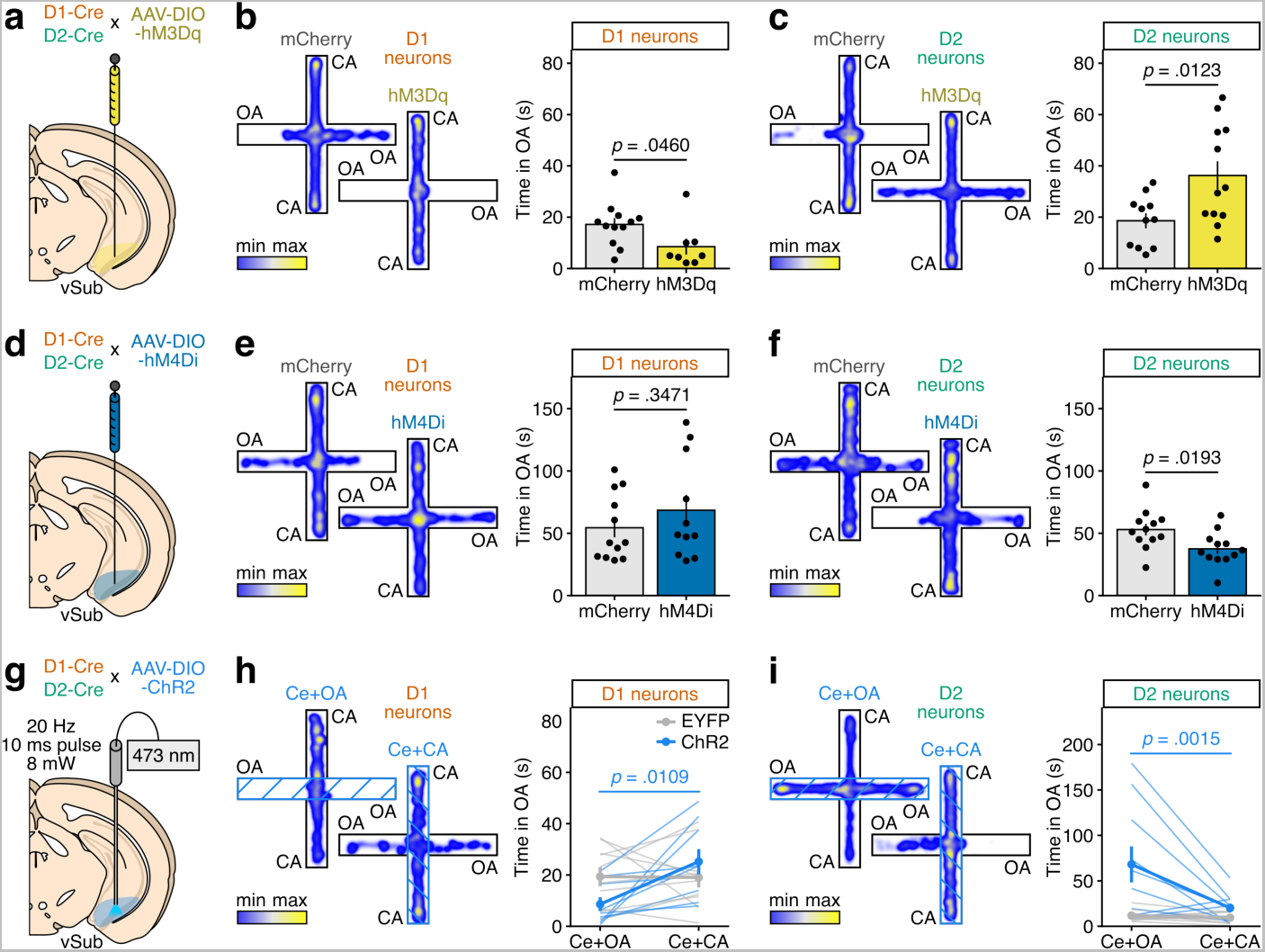
Chemogenetic and optogenetic manipulation of vSub D1 and D2 neurons during EPM testing. **a**, Experimental schematic. Male D1-Cre and D2-Cre mice were injected in vSub with an AAV-DIO-hM3Dq (n = 8 D1, n = 12 D2) or a control AAV-DIO-mCherry (n = 12 D1, n = 11 D2). CNO (3 mg/kg) was administered i.p. to all animals 15 min before testing. **b**, Representative examples of EPM exploration from D1-Cre mice (left) and quantification of open arm (OA) exploration time (right). Welch’s t-test: t_14.86_ = 2.1775 *p* = 0.0460. **c**, Representative examples of EPM exploration from D2-Cre mice (left) and quantification of open arm exploration time (right). Welch’s t-test: t_16.65_ = −2.8063 *p* = 0.0123. **d**, Experimental schematic. Male D1-Cre and D2-Cre mice were injected in vSub with an AAV-DIO-hM4Di (n = 11 D1, n = 12 D2) or a control AAV-DIO-mCherry (n = 12 D1, n = 12 D2). CNO (3 mg/kg) was administered i.p. to all animals 15 min before testing. **e**, Representative examples of EPM exploration from D1-Cre mice (left) and quantification of open arm exploration time (right). Welch’s t-test: t_16.97_ = −0.967 *p* = 0.3471. **f**, Representative examples of EPM exploration from D2-Cre mice (left) and quantification of open arm exploration time (right). Welch’s t-test: t_21.43_ = 2.5307 *p* = 0.0193. **g**, Experimental schematic. Male D1-Cre and D2-Cre mice were injected in vSub with an AAV-DIO-ChR2 (n = 9 D1, n = 10 D2) or a control AAV-DIO-EYFP (n = 10 D1, n = 11 D2). Optogenetic stimulation (473 nm laser, 8 mW, 20 Hz, 10 ms pulses) was delivered when the animal was either in the center or open arm (Ce+OA) or in the center and closed arm (Ce+CA) in a within-subject design. **h**, Representative examples of EPM exploration from D1-Cre mice (left) and quantification of open arm exploration time (right). LMM-ANOVA: stimulation zone F_1,17_ = 6.21 *p* = 0.0233, virus F_1,17_ = 0.26 *p* = 0.6145, stimulation zone x virus F_1,17_ = 6.72 *p* = 0.0190; followed by FDR-adjusted post-hoc tests. **i**, Representative examples of EPM exploration from D2-Cre mice (left) and quantification of open arm exploration time (right). LMM-ANOVA: stimulation zone F_1,19_ = 9.34 *p* = 0.0065, virus F_1,19_ = 8.96 *p* = 0.0075, stimulation zone x virus F_1,19_ = 7.68 *p* = 0.0121; followed by FDR-adjusted post-hoc tests. Data represented as mean ± sem.

To better mimic the spatiotemporal aspects of vSub D1 and D2 neuron activation at the time of approach/avoidance decision-making (Fig. 3), we turned to optogenetics, with stimulation of channelrhodopsin (ChR2)-expressing vSub D1 or D2 neurons (Fig. 4g) timed precisely to when the animal entered the EPM center or open arms (Ce+OA) and, as a within-subject control, when the animal entered the center or closed arms (Ce+CA). Real-time activation of D1 neurons in the open arm almost entirely abolished open arm exploration compared to closed arm stimulation in the same animals (Fig. 4h and Extended Data Figure 6h), while activation of D2 neurons dramatically increased open arm exploration time (Fig. 4i), number of entries and length of exploration bouts (Extended Data Fig. 6i) – effects absent in between-subject, EYFP-expressing controls. Interestingly, Ce+CA stimulation in D2 neurons increased the number of open arm entries but not their duration (Extended Data Fig. 6i), arguing against a purely place-preference effect and consistent with a pro-approach role for D2 neurons: D2 stimulation in the center platform might be enough to trigger open arm entry, which is then quickly cut short in the absence of continued D2 activation. Together, these experiments demonstrate that vSub D1 and D2 neurons oppositely modulate anxiety-related exploratory behaviors, and suggest that D2 activation during anxiogenic exploration promotes continuing the behavior, while the concomitant activation of D1 neurons dissuades it – consistent with correlative photometry data around head-dipping (Fig. 3f-j).

### Dopamine is released in vHipp during exploration of anxiogenic environments

These observations suggest that dopamine signals could differentially recruit either vSub D1 or D2 neurons and thus promote opposite approach/avoidance behaviors and exploratory outcomes. To determine the existence and spatiotemporal kinetics of dopamine release in vSub with behaviorally-relevant (second to sub-second) resolution, we took advantage of optical sensors for ultrafast *in vivo* dopamine imaging, which have yet to be tested in hippocampus where dopamine levels are relatively low and dynamics unknown ^44^. First, we used the D1-based sensor dLight-1.1 ^45^ and recorded photometry signals in vSub during EPM testing (Fig. 5). Albeit with poor signal quality that only allowed for the detection of gradual changes and not of unitary transients, dLight-1.1 signal intensity increased temporally and spatially with open arm exploration (Fig. 5b-e). Those changes were absent from control recordings obtained from animals expressing a non-dopamine-dependent GFP fluorophore (Fig. 5b,c,e and Extended Data Fig. 8f), and were recapitulated using either the D2-based sensor GRAB_DA_-1h ^46^ or red-shifted D1-based sensor RdLight-1 ^47^ – the latter offering improved *in vivo* dynamic range (Extended Data Fig. 8). Together, these data establish that anxiogenic exploration triggers dopamine release in vSub.

**Fig. 5:**
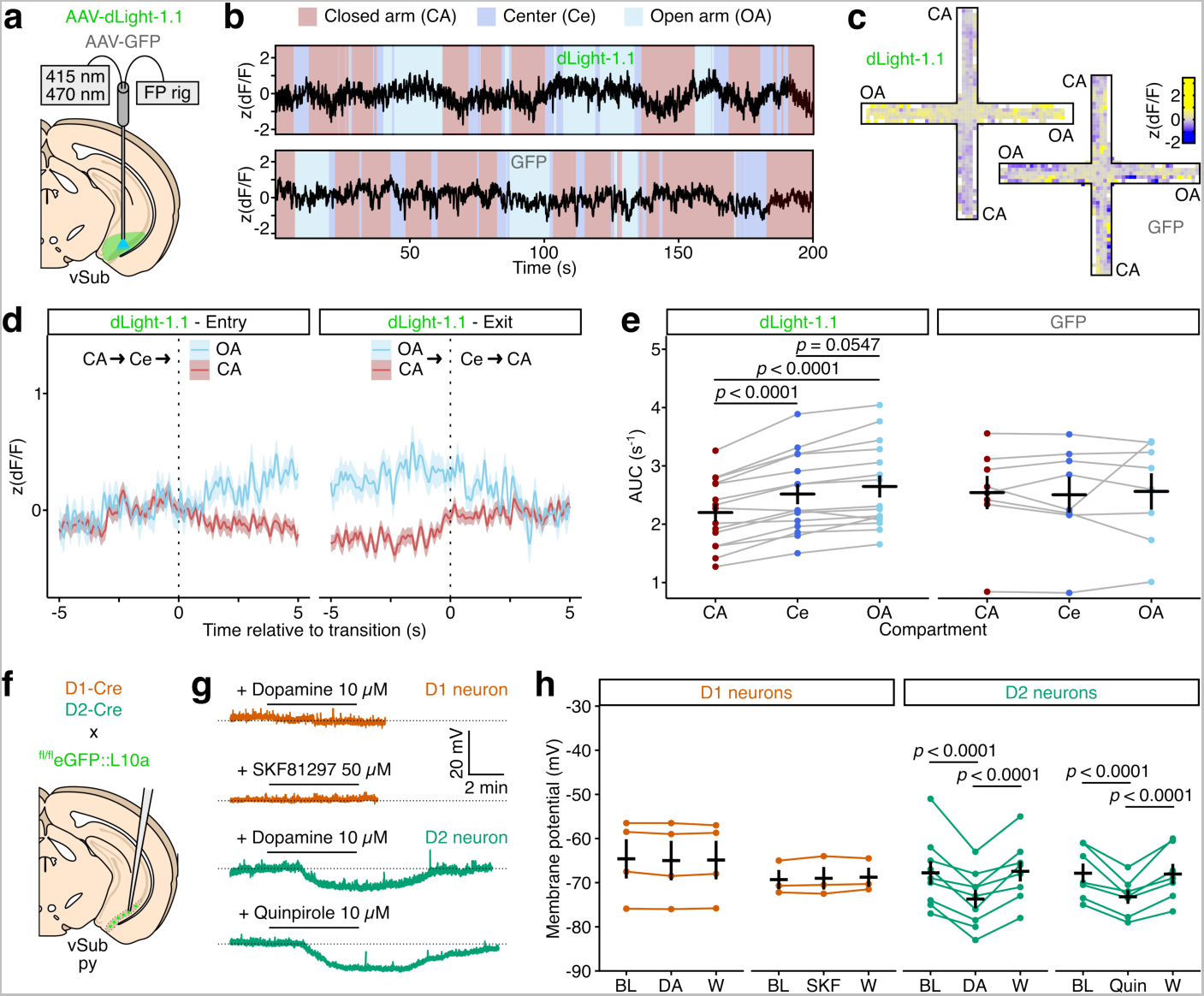
*In vivo* dopamine sensing and *ex vivo* dopamine pharmacology in vSub. **a**, Experimental schematic. Male mice were injected in vSub with an AAV-dLight-1.1 (n = 15), an AAV-GRAB_DA_-1h (n = 15, Extended Data Fig. 8), an AAV-RdLight-1 (n = 6, Extended Data Fig. 8) or with a control AAV-GFP (n = 8) and implanted with optical fibers before recording during EPM testing. **b**, Representative traces during EPM exploration for a dLight-1.1 (top) and control GFP (bottom) animal. **c**, Spatially averaged fluorescence signal in the EPM for all dLight-1.1 (top-left) and all GFP (bottom-right) mice. **d**, Average dLight-1.1 signal during entries to (left) and exits from (right) the open arm (OA, blue) or closed arm as a control (CA, red). Only traces when the mouse did a complete closed arm to center (Ce) to open/closed arm are used for averaging. **e**, Average area under the curve (AUC) by EPM compartment for dLight-1.1 (left) and GFP (right) mice. LMM-ANOVA: compartment F_2,80_ = 49.82 *p* < 0.0001, sensor F_3,40_ = 1.70 *p* = 0.1822, compartment x sensor F_6,80_ = 5.1529 *p* = 0.0002; followed by FDR-adjusted post-hoc tests. **f**, Experimental schematic for slice electrophysiological recordings from vCA1/vSub D1 or D2 pyramidal neurons identified in D1-Cre or D2-Cre x ^fl/fl^eGFP::L10a male mice. **g**, Representative traces of resting membrane potential before, during and after bath application of dopamine (DA, 10 µM) or of the D1 agonist SKF81297 (SKF, 50 µM) onto D1 neurons (top), or of dopamine (DA, 10 µM) or of the D2 agonist quinpirole (Quin, 10 µM) onto D2 neurons (bottom). **h**, Quantification of resting membrane potential changes between baseline (BL), drug application and after wash (W). LMM-ANOVA: cell type F_1,5.8_ = 0.90 *p* = 0.3805, agonist F_1,18.7_ = 0.40 *p* = 0.5366, application period F_2,37.0_ = 17.73 *p* < 0.0001, cell type x agonist F_1,18.7_ = 0.51 *p* = 0.4851, cell type x application period F_2,37.0_ = 16.39 *p* < 0.0001, agonist x application period F_2,37.0_ = 0.23 *p* = 0.7972, cell type x agonist x application period F_2,37.0_ = 0.02 *p* = 0.9821; followed by FDR-adjusted post-hoc tests. Data represented as mean ± sem.

### vSub D1 and D2 neurons differently respond to dopamine

We next set out to check the postsynaptic consequences of dopamine release onto vSub D1 and D2 neuronal activity. In striatum, dopamine binding to a D1 receptor – coupled to Gα_s_ – favors activation of D1-expressing neurons, while dopamine binding to a D2 receptor – coupled to Gα_i/o_ – exerts an inhibitory effect on D2-expressing neurons ^48^. We performed cell-type-specific *ex vivo* slice electrophysiological recordings of vCA1/vSub D1 and D2 pyramidal neurons identified in the same transgenic mouse lines used above (Fig. 5f). In current-clamp mode, robust yet reversible hyperpolarization was observed in D2 neurons after application of either dopamine itself or of the D2 agonist quinpirole (Fig. 5g,h). However, neither dopamine nor the D1 agonist SKF81297 caused depolarization of D1 neurons at the concentrations tested (Fig. 5g,h), suggesting more subtle excitatory effects of dopamine on this cell type. In any case, these findings support earlier conclusions that vCA1/vSub D1 and D2 neurons represent distinct and non-overlapping populations (Figs. 1,2).

We propose that dopamine acts as a “bottom-up” signal onto vCA1/vSub D1 and D2 neurons to gate exploratory behaviors in anxiogenic environments by arbitrating the further recruitment of one or the other cell type. As such, vCA1/vSub dopamine levels might operate as a threshold to limit further investigation under anxiety-like conditions if/when they rise enough to shift the ratio of D1-/D2-mediated signaling. Importantly, our findings do not imply that dopamine is the main driver of these two dopaminoceptive neuronal subpopulations – which presumably result from other, both intra- and extra-hippocampal, glutamatergic innervation ^6,7,13,14^ – but rather reflect dopamine’s general role as a neuromodulator, fine-tuning the integration of these input signals.

### vSub dopamine signaling contributes to conditioned approach and active avoidance

Finally, we tested whether this model extends to more sophisticated measures of approach/avoidance decision-making. To that end, we adapted for mice a platform-mediated avoidance (PMA) task previously developed for rats ^49–51^ to interrogate the vHipp dopamine correlates of active avoidance *versus* motivated foraging/approach strategies elicited by learned, discrete, threat-predicting cues (Fig. 6a, see Methods for detailed description). As such, this task transforms the non-descript, spatially-determined conflict present in EPM into a temporally-precise one with better-defined motivational dimensions, and it incorporates elements of novelty and learning particularly relevant when studying hippocampal and dopamine systems ^52,53^. First, we recorded photometry signals for D1 and D2 activity (cell-type-specific GCaMP6s expression in D1- or D2-Cre male mice) or dopamine release (RdLight-1 probes) in vSub during a first PMA conditioning session. Electric foot-shocks robustly triggered D1 and D2 activity, as well as dopamine release (Fig. 6b left and Extended Data Fig. 9a left) – adding vSub to the list of brain regions receiving aversion-related dopamine signals ^10,31^. Reward consumption also elicited dopamine release, which was accompanied by a decrease in D2 activity (Fig. 6b middle and Extended Data Fig. 9a middle), recapitulating canonical aspects of dopamine signals in reward learning ^27,30^. No increase in D1 activity was detected, suggesting that dopamine concentrations might in this case reach D2 but not D1 receptor affinity levels – D2 receptors have 10-100-fold higher affinity for dopamine *in vitro* ^54^. To infer approach/avoidance conflict resolution, photometry signals were analyzed around the first platform-to-grid exit transition immediately after the tone ends (Fig. 6b right and Extended Data Fig. 9a right), when we expect the uncertainty between safety and exploration to dictate decision-making. D2 neurons activity peaked ±2.5 s before the mouse exited the platform – consistent with the pro-approach, motivating role hypothesized above for D2 cells – and this transition occurred concurrently with a sustained increase in dopamine release. Moreover, for both foot-shocks and platform transition, dopamine activity rose after the peak of D2 activity (Fig. 6c and Extended Data Fig. 9a), and for all events changes in dopamine levels and D2 activity were negatively correlated with dual GCaMP6s/RdLight-1 expression (Extended Data Fig. 9b), consistent with the idea that dopamine acts to turn off D2 cells in these timeframes. Other control, time-locked behavioral events (*e.g.*, active lever presses, tone onset, platform mounting) did not elicit any response in either D1, D2 or dopamine signals (data not shown).

**Fig. 6:**
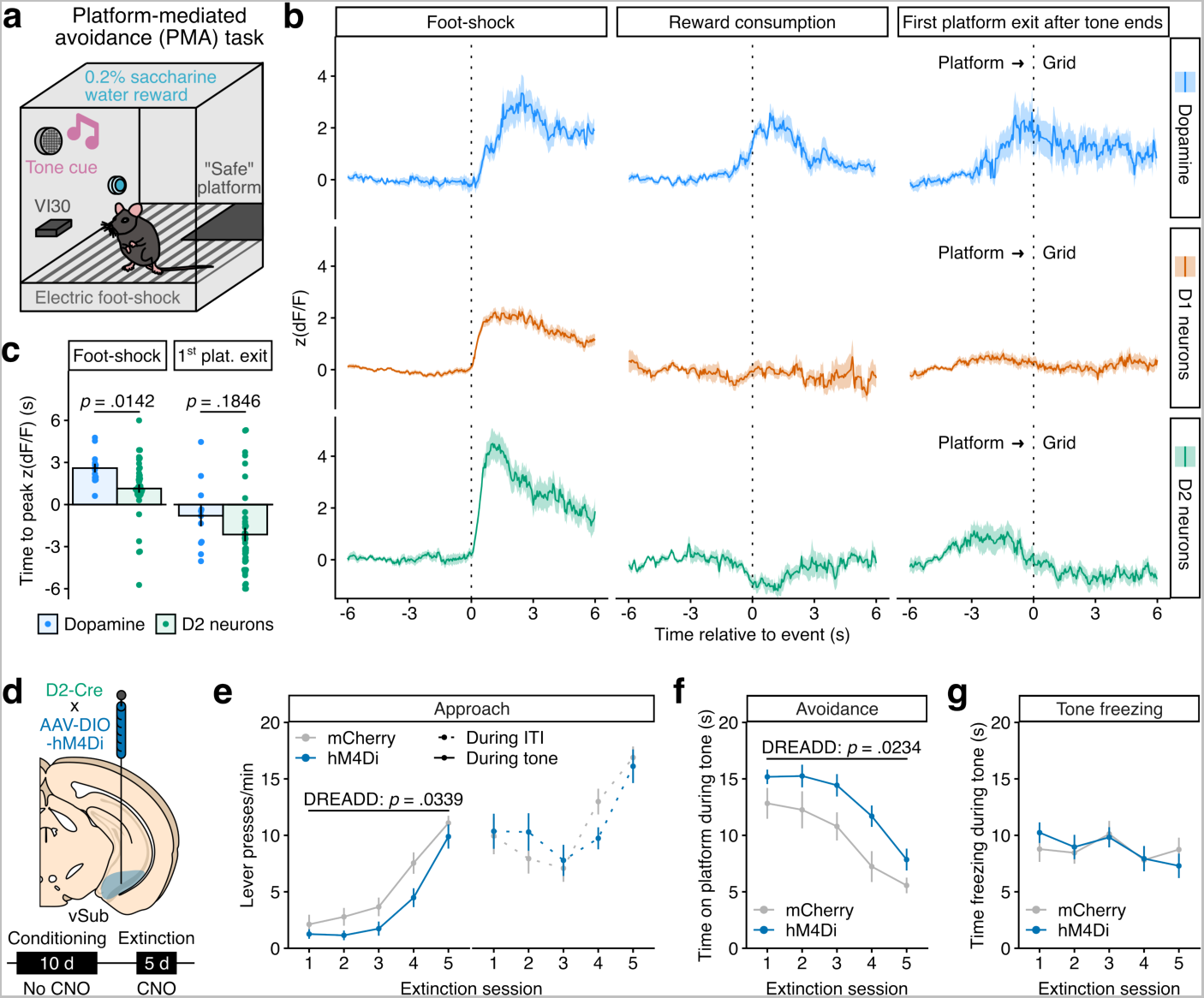
vSub dopamine, D1 and D2 correlates of approach and avoidance in the PMA task. **a**, Experimental schematic of the PMA task (see Methods for detailed description). **b**, Fiber photometry recordings in male mice injected in vSub with an AAV-RdLight-1 (top, n = 2), or D1-Cre (middle, n = 13) and D2-Cre (bottom, n = 8) mice injected with an AAV-DIO-GCaMP6s and implanted with optical fibers before recording in the PMA task. Average signal traces are centered around the onset of the electric foot-shock, when the mouse is on the grid (left), the entry into the reward magazine for reward consumption (middle) or the exit from the platform towards the grid, the first time after the tone and shock end if the mouse was on the platform (right). Quantification and statistics in Extended Data Fig. 9. **c**, Times of peak photometry RdLight-1 and D2-GCaMP6s signals after foot-shock (left, LMM-ANOVA: sensor F_1_ = 6.41 *p* = 0.0142) and first platform exit (right, LMM-ANOVA: sensor F_1_ = 1.81 *p* = 0.1846). **d**, Experimental schematic. Male D2-Cre mice were injected in vSub with an AAV-DIO-hM4Di (n = 9) or a control AAV-DIO-mCherry (n = 11). PMA training was run for 10 days without CNO treatment, then CNO (3 mg/kg) was administered i.p. to all animals 15 min before testing under extinction conditions (no shock) for 5 days. **e**, Approach behavior (lever presses) during extinction sessions, both during tone presentation (left; LMM-ANOVA: session F_4,72_ = 64.75 *p* < 0.0001, DREADD F_1,18_ = 5.27 *p* = 0.0339, session x DREADD F_4,72_ = 0.83 *p* = 0.5099) and inter-trial intervals (ITI, right; LMM-ANOVA: session F_4,72_ = 22.15 *p* < 0.0001, DREADD F_1,18_ = 0.006 *p* = 0.9379, session x DREADD F_4,72_ = 2.03 *p* = 0.0994). **f**, Avoidance behavior (time on platform) during extinction sessions (LMM-ANOVA: session F_4,72_ = 26.69, *p* < 0.0001, DREADD F_1,18_ = 6.13 *p* = 0.0234, session x DREADD F_4,72_ = 0.57 *p* = 0.6858). **g**, Freezing upon tone presentation during extinction sessions (LMM-ANOVA: session F_4,72_ = 4.25 *p* = 0.0038, DREADD F_1,18_ = 0.003 *p* = 0.9570, session x DREADD F_4,72_ = 1.45 *p* = 0.2278). Data represented as mean ± sem.

To causally link the increase in D2 vSub neuron activity to approach behavior in the PMA task, we employed D2-specific inhibitory chemogenetics (Fig. 6d), which produced the most convincing behavioral effects in EPM (Fig. 4f) and novelty-suppressed feeding (Extended Data Fig. 7d). CNO was given during extinction sessions, when mice have to engage in approach, exploratory behavior during tone presentation to update punishment contingencies and maximize reward consumption. Both hM4Di- and control mCherry-expressing mice readily acquired approach and avoidance strategies (Extended Data Fig. 10). Under extinction conditions, however, inhibition of D2 neurons decreased the rate of lever pressing (Fig. 6e left) and increased the time spent on the ‘safe’ platform (Fig. 6f) during tone presentation, indicating decreased approach and increased avoidance. Importantly, this manipulation did not affect lever pressing behavior during inter-tone intervals (Fig. 6e right), denoting that these effects might be selective to conflicting situations. D2 neurons’ inhibition also did not affect freezing behavior (Fig. 6g), suggesting specificity to active *versus* passive avoidance. While these effects could be interpreted as general deficits in extinction learning, we argue that they arise from reduced motivation to engage in exploratory behavior during signaled threat prediction.

## Discussion

Our findings strongly support the existence of an anatomical and functional dichotomy between vHipp D1 and D2 neurons in vSub, with segregated D1 and D2 neurons cooperating to balance distinct and in some cases opposite behaviors related to motivated exploration and investigation in the face of anxiety on one hand, and anxiety-like features detection and avoidance on the other (Fig. 4) – in a manner compellingly reminiscent of extensively-studied D1 and D2 striatal MSNs ^26–29^ and where dopamine signals (Fig. 5) might similarly operate to shift the relative recruitment of those two subpopulations during decision-making ^30^. One key aspect resides in the precise circuit identity of those vCA1/vSub D1 and D2 pyramidal neurons, in terms of both the inputs they receive (as they exhibit partially distinct activation patterns, Figs. 3,6) and their target projections (as they promote distinct outcomes, Figs. 4,6), especially given that subtle input-output biases have been reported in this region ^13,14^. Local connections with D1 or D2 interneurons in vHipp, as well as dendritic-compartment-specific dopamine receptor expression ^55^ and/or layer-specific dopamine innervation ^19,21^, could create additional heterogeneity of dopamine-processing functional units – which might be encapsulated by shared transcriptional features (Fig. 2). Moreover, with vCA1/vSub projecting to the prefrontal cortex and nucleus accumbens ^13,14^ – which in turn projects back to the midbrain – dopamine axons in vHipp form functional polysynaptic loops, which are hypothesized to be crucial for long-term memory formation ^52^. Hence, dopaminoceptive vCA1/vSub cells might constitute a sub-group of hippocampal neurons that are especially well integrated within the mesocorticolimbic dopamine circuitry and participate in dopamine signal processing both locally in vHipp and at the nerve-terminal level in the nucleus accumbens ^56^. The ascending arc of those loops implicates midbrain-originating innervation of vCA1/vSub, which has been observed experimentally ^18–20^. Given the diversity of midbrain dopamine neurons and their limited collateralization ^57^, it likely arises from unique populations with idiosyncratic intrinsic properties, which might explain the distinctive dopamine release profiles we observed here (Figs. 5,6). Another – provocative – hypothesis derives from recent studies of dopamine signaling in dorsal hippocampus, where a significant, if not preponderant, portion of dopamine there is co-released from noradrenergic fibers from the locus coeruleus ^58^, with separate roles related to novelty detection and memory encoding for midbrain- *versus* locus coeruleus-originating dopamine ^53^. Whether this is also true in vHipp is currently unknown, but it typifies the ever-growing complexity of the diverse neuromodulatory inputs that converge into vHipp: dopamine in all likelihood acts in concert with norepinephrine, serotonin, acetylcholine and other modulatory molecules (including peptides) to dictate appropriate behavioral selection under intertwined novelty, anxiety and memory-encoding conditions. From a translational standpoint, delineating the precise neuromodulatory circuits that govern anxiety is an essential step forward in addressing this leading cause of disability worldwide.

## Methods

### Animals

Male C57BL/6J mice (8-32 weeks old, 20-30 g, The Jackson Laboratory) were maintained on a 12:12 h light/dark cycle (07:00 lights on; 19:00 lights off) and were provided with food and water *ad libitum*. Transgenic mouse lines (D1-Cre: MGI:3836633, D2-Cre: MGI:3836635, D1-tdTomato: MGI:4360387, ^fl/fl^eGFP::L10a: IMSR_JAX:022367) were bred in-house on a C57BL/6J background. All mice were maintained according to the National Institutes of Health guidelines for Association for Assessment and Accreditation of Laboratory Animal Care accredited facilities. All experimental protocols were approved by the Institutional Animal Care and Use Committee at Mount Sinai. All anxiety-related behavioral testing was done between 2 and 6 h into the dark phase, while the PMA task was done during the light phase.

### Drug treatments

CNO (Tocris) was first diluted in DMSO (Sigma), then in Dulbecco’s Phosphate Buffered Saline (PBS, Gibco) to a 0.5% DMSO final concentration and injected intraperitoneally at 3 mg/kg 15 min before behavioral testing. Dopamine hydrochloride (Tocris, #3548), SKF81297 hydrobromide (Tocris, #1447) and (-)-Quinpirole hydrochloride (Tocris, #1061) were diluted in ACSF (see below) to respectively 10, 50 and 10 µM, extemporaneously for each day of recording, and kept protected from light for the duration of the bath perfusion.

### Viral reagents

The following viruses were obtained from Addgene: for chemogenetics, AAV9-hSyn-DIO-hM3D(Gq)-mCherry (#44361), AAV9-hSyn-DIO-hM4D(Gi)-mCherry (#44362), AAV9-hSyn-DIO-mCherry (#50459); for optogenetics, AAV9-EF1a-DIO-hChR2(H134R)EYFP-WPRE-HGHpA (#20298), AAV9-EF1a-DIO-EYFP (#27056); for calcium imaging, AAV9-CAG-Flex-GCaMP6s-WPRE-SV40 (#100842); for dopamine sensing, AAV5-CAG-dLight1.1 (#111067), AAV9-hSyn-GRAB_DA1h (#113050), AAV9-CMV-PI-EGFP-WPRE-bGH (#105530). For red-shifted dopamine sensing, the AAV-DJ/2-CAG-RdLight1-WPRE-SV40 was obtained from the Viral Vector Facility of the ETH Zürich (#v577-DJ). All viruses were used at ±1 × 10^12^ GC/mL, except AAV9-CAG-Flex-GCaMP6s-WPRE-SV40 at ±5 × 10^12^ GC/mL, AAV5-CAG-dLight1.1 and AAV9-hSyn-GRAB_DA1h at ±2 × 10^13^ GC/mL.

### Stereotaxic surgeries

Mice were anesthetized with an intraperitoneal bolus of ketamine (100 mg/kg) and xylazine (10 mg/kg), then head-fixed in a stereotaxic apparatus (Kopf Instruments). Syringe needles (33G, Hamilton) were used to bilaterally infuse 1 µl of virus at a 0.1 µl/min flow rate, except for fiber photometry experiments where infusion was unilateral. Needles were kept in place for 10 minutes after injection before being retracted to allow for virus diffusion. Coordinates for vSub were as follows, from bregma: AP –3.3 mm, ML +2.9 mm, DV –4.5 mm, 0° angle. For fiber photometry, 400 µm-wide optical fibers (Doric, MFC_400/430-0.66_4.5mm_MF2.5_FLT) were unilaterally implanted at AP –3.3 mm, ML +2.9 mm, DV –4.4 mm, 0° angle. For optogenetics, 200 µm-wide optical fibers (Doric, MFC_200/240-0.22_4.5mm_MF1.25_FLT) were bilaterally implanted above vSub at AP –3.3 mm, ML +2.9 mm, DV –4.3 mm, 0° angle. Optical fibers were secured in place using dental cement (3M) without the use of screws to the skull and covered with a layer of black dental cement (C&B Metabond). Virus infusion and optical fiber placement was confirmed either by immunohistochemistry on fixed brains sections or by dissection of fresh tissue under fluorescent light. Representative examples of viral expression and fiber placement are shown in Extended Data Fig. 3.

### Immunohistochemistry and imaging

Mice were transcardially perfused with a fixative solution containing 4% paraformaldehyde (PFA). Brains were post-fixed for 24 h in 4% PFA at 4°C. Sections of 40 µm thickness were cut in the coronal plane with a vibratome (Leica) and stored at −20°C in a cryoprotectant solution containing 30% ethylene glycol (v/v), 30% glycerol (v/v) and 0.1 M phosphate buffer. Native GFP signals in D1- or D2-Cre x ^fl/fl^eGFP::L10a mice were enhanced by immunohistochemistry using an anti-GFP primary antibody (chicken; 1:500; #GFP-1020, Aves Labs). Sections were finally incubated with secondary antibodies (donkey anti-chicken Alexa-488-conjugated (1:500; Jackson Immunoresearch), counterstained with DAPI and mounted in ProLong Gold Antifade Mountant (ThermoFisher Scientific). For viral placement verification (Extended Data Fig. 3), no immunohistochemistry was performed. Confocal images (1024 × 1024 pixels, 16 bits pixel depth, pixel size: x = 1.14 µm, y = 1.14 µm, z = 2.6 µm) were acquired on a SP8 inverted confocal microscope (Leica) using a 10X objective and Leica Application Suite x v3.5.7.23225. Entire hemispheres images (6392 × 5344 µm) were reconstructed from 30 images (1163.64 × 1163.64 µm) stitched using ImageJ software.

### Nuclei purification and fluorescence-activated nuclei sorting (FANS)

Mouse brains were collected after cervical dislocation and followed by rapid bilateral vHipp dissections from 1 mm-thick coronal brain sections using a 14G needle and frozen on dry ice. To obtain a nuclei suspension, frozen vHipp samples were homogenized in 4 mL of low-sucrose lysis buffer (0.32 M sucrose, 5 mM CaCl_2_, 3 mM Mg(Ace)_2_, 0.1 mM EDTA, 10 mM Tris-HCl) using a large clearance then a small clearance pestle of a glass dounce tissue grinder (Kimble Kontes). Homogenates were filtered through a 40 µm cell strainer (Pluriselect) into ultracentrifuge tubes (Beckman Coulter), underlaid with 5 mL of high-sucrose solution (1.8 M sucrose, 3 mM Mg(Ace)_2_, 1 mM DTT, 10 mM Tris-HCl) and centrifugated at 107,000 *g* for 1 hour at 4°C in a SW41Ti Swinging-Bucket Rotor (Beckman Coulter). Supernatant was discarded and nuclei pellets were re-suspended in 800 µL of PBS supplemented with 0.5% bovine serum albumin (BSA). DAPI was added at a 1 µg/mL final concentration. Nuclei were sorted on a BD FACS Aria II three-laser device with a 70 µm nozzle and using BD FACSDiva Software v8.0.2. Gating strategy from a representative sort are visualized in fig. S1. Briefly, debris and doublets were excluded using FSC and SSC filters, nuclei were then selected as DAPI-positive (Violet1-A laser) events, and finally GFP-positive nuclei (Blue1-A laser) were sorted directly into BSA-coated low-binding tubes. 25,000 nuclei were recovered for each sample.

### Single-nuclei RNA-sequencing (snRNAseq) and analysis

Following FANS, nuclei were quantified (Countess II, Life Technologies) and 8,000 per sample were loaded on a single 10X lane using Chromium Single Cell 3’ Library Construction Kit (10X Genomics). cDNA libraries were prepared according to the manufacturer’s protocol (10X Genomics). Libraries were sequenced at New York Genome Center using the NovaSeq platform (Illumina) at a depth of ±150 millions reads per sample. A Cell Ranger (v3.1.0) reference package was generated from the mm10 pre-mRNA mouse genome that ensured alignment to unspliced pre-mRNAs and mature RNAs. Cell Ranger filtered outputs were analyzed with *Seurat v4.3.0* in R v4.2.2. Nuclei containing <900 reads, or <200 or >5000 features (*i.e.,* genes for which at least one read was detected), or >1% of reads mapping to the mitochondrial genome were removed, leaving respectively 11,452 and 18,158 nuclei from D1 and D2 samples for further analysis. Nuclei from all samples then underwent integration using 3,000 features for *FindIntegrationAnchors*, clustering using 32 principal components and 40 nearest neighbors for *FindNeighbors* and a 0.5 resolution value for *FindClusters* following *Seurat v4.3.0* vignette. These values were determined to recapitulate previously defined cell types. UMAP dimensionality reduction was finally run with *RunUMAP* on the *integrated_snn* graph calling the *r-reticulate* Python v3.6.10 install of *umap-learn v0.4.6* for visualization purposes. Marker genes for each cluster were computed with *FindAllMarkers* and regressing out sample identity using logistic regression (full marker gene lists and statistics available in Supplementary Information Table 1). Individual clusters were then further manually annotated by comparing enriched marker genes for each cluster (Extended Data Fig. 2) with publicly available single-cell RNAseq databases of whole brain tissue ^36^, hippocampal ^37,38^ and subicular ^39^ principal cells, and hippocampal interneurons ^40^. One cluster (pyramidal neurons [PNs] #2) was only enriched for one gene (*Gm41418*, associated with ribosomal RNA contamination) and comprised nuclei with lower than average read counts, and was thus labeled as “low quality (_LQ_)”. Cluster-specific differential expression analysis between D1 and D2 nuclei was performed using *FindMarkers*. Fold change was computed for all genes, but statistical testing using logistic regression was further restricted only to genes detected in >25% of either D1 or D2 nuclei in each cluster, and with >20% expression change between D1 and D2 nuclei. *p*-values were adjusted for false discovery rate (FDR) at a 0.05 significance level (full DEG lists and statistics available in Supplementary Information Table 2). Hierarchical clustering of DEGs was then run on union heatmaps comprising genes that were significantly regulated (FDR-adjusted *p* < 0.05) in at least one cell type cluster, separately for GABAergic and glutamatergic cells, with the *cluster v2.1.4::agnes* function in R v4.2.2 using Ward’s minimum variance and computing Euclidian distances (full pattern gene lists and statistics available in Supplementary Information Table 3). Gene ontology (GO) enrichment analyses of gene expression patterns was then performed using the PANTHER knowledgebase and classification system ^59^ (full GO term lists and statistics available in Supplementary Information Table 4).

### Fiber photometry recordings and analysis

To analyze bulk photometry signals from GCaMP6s or dopamine-sensing probes during mouse behavior, the fiber photometry system was time-locked with the video-tracking system (Ethovision XT 11, Noldus or ANY-Maze) or the MedPC operant systems via transistor–transistor logic signals (TTLs). Two different photometry systems were used; The first ^11,60^ was used for initial dLight-1.1 and GRAB_DA_-1h recordings, and used two light-emitting diodes at 490 and 405 nm (Thorlabs), reflected off dichroic mirrors (FF495; Semrock) and coupled to optical fibers using a 6 m-long low-autofluorescence patchcord (Doric, MFP_400/430/1100-0.66_3m_FCM-MF2.5_LAF). The real-time fiber photometry signal was collected using a signal processor (Tucker–Davis Technologies, TDT) and acquired with open source OpenEx software 2.20 controlling an RX8 lock-in amplifier (Tucker-Davis Technologies). OpenEx, sinusoidally modulated each LED’s output (490 nm at 211 Hz, and 405 nm isosbestic control at 531 Hz). The two output signals were then projected onto a photodetector (2151 fW photoreceiver; Newport). The photoreceiver signal was sampled at 6.1 kHz, after which each of the two modulated signals was separated by the real-time processor for analysis. Decimated signals were collected at a sampling frequency of 381 Hz to perform the post-acquisition analyses. The second (FP3002 system from NeuroPhotometrics, NPM) had dual color capability and was used for later dLight-1.1 and GRAB_DA_-1h recordings, as well as for all GCaMP6s and RdLight-1 recordings. It was similarly connected to optical fiber head implants using the same low-autofluorescence patchcords, and was used according to manufacturer’s instructions and with the FP3002 Bonsai node. Fluorescence signals resulting from 470 nm, 560 nm and 415 nm excitation, interleaved in time, were sampled at 78 Hz total, *i.e.* at a 26 Hz effective sampling rate for each channel.

Post-acquisition analyses were performed using custom programs similarly for data obtained from both set ups. First, TDT data were extracted and converted into fluorescence time-series using generic MATLAB code from the Lerner lab (https://github.com/talialerner/). Deinterleaved NeuroPhotometrics time series data was directly exported from Bonsai. To compare neuronal activity across animals and behavioral sessions, individual animal time-series data were analyzed using custom R codes following published standard methods ^61^ with minor modifications. Briefly, both 490 and isosbestic 405 nm (for TDT) or 470, 560 and isosbestic 415 nm (for NPM) signals were first smoothed using a 4th-order 5 Hz lowpass Butterworth filter built using the *gsignal v0.3-5::butter* function. To remove the bleaching slope and low-frequency fluctuations, baseline correction was then performed by subtracting the baseline obtained by regressing each individual signal using the LOWESS smoother (*stats::lowess*) with default parameters from the smoothed 490/470/560 and 405/415 nm signals. Both 490/470/560 and 405/415 nm signals were then standardized using a robust z-score (zF = (F – median(F))/mad(F)). The standardized 405/415 nm signal was then fitted to the standardized 490 nm signal using the robust regression function *MASS v7.3::rlm*, and normalized dF/F z(dF/F) was finally calculated as the difference between the 490/470/560 nm signal and the fitted 405/415 nm signal to remove motion artifacts and autofluorescence. To analyze time-locked neuronal activity in respect to behavior, the normalized z(dF/F) signal was extracted around the onset of the relevant behavior (defined as t = 0 s). For compartment-based spatial analysis, signal changes were quantified for relevant time intervals (in each compartment) as the corresponding areas under the curve (the curve being the entire z(dF/F) time-series), which were calculated with linear interpolation using the *MESS v0.5.9::auc* function. For peri-event analysis, signal trace data are quantified and averaged with n = event, and signal changes are measured as the maximum (peak) value reached by each z(dF/F) time-series within relevant time intervals. Signal change slope was calculated as the average of first derivative slope values of each z(dF/F) time-series in the 200 ms window around the time of maximal slope.

Unsupervised hierarchical clustering was run on D1 and D2 time-series together with the *cluster v2.1.4::agnes* function in R v4.2.2 using Ward’s minimum variance and computing Euclidian distances. Supervised, non-probabilistic binary linear classification using support vector machines (SVM) was then run separately on D1 and D2 time-series using the entire head-dip-centered time-series as input (26 Hz sampling for 3 s = 78 input features). Datasets were first randomly split into training (75%) and testing (25%) sets. SVM model training was achieved on the training set using C-type classification, a linear kernel and 5-fold cross validation with cost tuning in R v4.2.2 using the *e1071 v1.7::tune.svm* function. The trained model was then presented with testing set data, and SVM-predicted outcomes were compared to testing set observed outcomes to compute accuracy. The operation was repeated 10 times with a new random split of training/testing sets to perform statistical comparisons against chance performance (50% decoding accuracy) and between D1 and D2 signals.

### Ex vivo slice electrophysiology

Mice (12-20 weeks old) were deeply anesthetized with isoflurane and decapitated. Brains were rapidly removed and chilled in artificial cerebrospinal fluid (ACSF) containing (in mM): N-methyl-D-glucamine 93, HCl 93, KCl 2.5, NaH_2_PO_4_ 1.2, NaHCO_3_ 30, HEPES 20, glucose 25, sodium ascorbate 5, thiourea 2, sodium pyruvate 3, MgSO_4_ 10, and CaCl_2_ 0.5, pH 7.4. The brain was embedded in 2% agarose and coronal slices (200 µm thick) were made using a Compresstome (Precisionary Instruments). Brain slices were allowed to recover at 33 ±1°C in ACSF solution for 30 min and thereafter at room temperature in holding ACSF, containing (in mM): NaCl 92, KCl 2.5, NaH_2_PO_4_ 1.2, NaHCO_3_ 30, HEPES 20, glucose 25, sodium ascorbate 5, thiourea 2, sodium pyruvate 3, MgSO_4_, and CaCl_2_ 2, pH 7.4. After at least 1 h of recovery, the slices were transferred to a submersion recording chamber and continuously perfused (2-4 mL/min) with ACSF containing (in mM): NaCl 124, KCl 2.5, NaH_2_PO_4_ 1.2, NaHCO_3_ 24, HEPES 5, glucose 12.5, MgSO_4_ 2, and CaCl_2_ 2, pH 7.4. All the solutions were continuously bubbled with 95% O_2_ / 5% CO_2_. vCA1 or vSub pyramidal cells were visually identified with infrared differential contrast optics (BX51; Olympus) and fluorescence visualized through eGFP bandpass filters upon LED illumination through the objective (p3000^ULTRA^, CoolLed) using µManager v2.0. Whole-cell patch-clamp recordings were performed at room temperature using a Multiclamp 700A amplifier (Molecular Devices). Recording electrodes (3-5 MΩ) pulled from borosilicate glass were filled with solution containing (in mM): K-gluconate 122, HEPES 10, KCl 5, MgATP 5, Na_2_GTP 0.5, QX314 1, and EGTA 1, pH 7.25. Data acquisition (filtered at 10 kHz and digitized at 10 kHz) and analysis were performed with pClamp 11 software (Molecular Devices). Following the breakthrough cells were allowed to stabilize for 3 min before the recordings were made. Neurons were current clamped at I = 0 and baseline membrane potentials were recorded. Membrane potential values were corrected for liquid junction potential, which was determined empirically.

### Elevated-plus maze (EPM)

Two EPM apparatuses were used, explaining differences in baseline time in open arms in controls groups between cohorts. The first was designed in black Plexiglass (arm length 70 cm, arm width 5 cm, height 80 cm) and fitted with white surfaces to provide contrast. The second was similar but had wider arms (arm length 75 cm, arm width 8 cm, height 80 cm). Testing conditions occurred under red-light conditions (10 lux) in a room isolated from external sound sources. The EPM apparatus was thoroughly hand cleaned between mice with an odorless 30% ethanol cleaning solution. Mice were positioned in a closed arm, and behavior was video tracked for a 5 min period. Time in EPM compartments, locomotion, and velocity were measured using a video tracking system (Ethovision) set to localize the mouse center point at each time of the trial. Head-dips were time-stamped and the corresponding outcomes (avoid/explore/unclear) manually classified off-line by an experimenter blind to the groups.

### Open field test (OFT)

Mice were placed in the open field arena (44 × 44 cm) for 5 min to compare the distance traveled and time spent in the peripheral zone (7.5 cm from each border wall) compared to the center zone. Testing conditions occurred under red-light conditions (10 lux) in a room isolated from external sound sources. The OFT apparatus was thoroughly hand cleaned between mice with an odorless 30% ethanol cleaning solution. The mouse’s activity – distance, velocity, and time spent in specific open field areas – was measured for a 10 min period using a video tracking system (Ethovision) set to localize the mouse center point at each time of the trial.

### Novelty-suppressed feeding (NSF) test

Mice were food-restricted for 24 h before testing occurred. Mice were then placed in the corner of a novel, black open-field arena (44 × 44 cm) covered in a different type of saw-dust bedding and where a single pellet of chow food was placed in the center of the arena. Testing conditions occurred under red-light conditions (10 lux) in a room isolated from external sound sources. The NSF boxes were thoroughly hand cleaned between mice with an odorless 30% ethanol cleaning solution, bedding replaced and a new food pellet was used for each mouse. Latency to feed was hand-scored by an experimenter blind to the experimental groups as the start of the first bout of uninterrupted feeding that was longer than 5 s.

### Optogenetics

For optogenetic stimulation experiments, mice were connected to a dual optical fiber patchcord (Doric) connected to a 473 nm blue laser (OEM Laser System). Stimulation was executed in the form of 10 ms box pulses emitted at 20 Hz with an output power of ±8 mW at the tip of the fiber. Optogenetic stimulation was triggered based on the mouse center-point location in the EPM tracked using Ethovision XT 11 (Noldus): for the first 5 min, when the mouse was in the center or open arm of the maze (Ce+OA stimulation), for the last 5 min, when the mouse was in the center or closed arm of the maze (Ce+CA stimulation).

### Platform-mediated avoidance (PMA) task

The PMA task consists of three phases: pre-training, conditioning, and extinction, carried out in Med Associates operant chambers. During the pre-training phase (3-7 days), water-restricted mice (2 mL of daily intake) learned to press a lever for a saccharine-water reward (0.2% saccharine). The mice first learned to press under a fixed-ratio 1 (FR1) reward-delivery schedule, which we increasingly delayed to variable interval (VI) 10, then VI20 and finally VI30 to criterion (>10 lever presses/min). Next, during the conditioning phase (10 days), the mice were exposed to 9 tones (20 s, 4 kHz, 75 dB) co-terminated with a 2-sec foot-shock (0.4 mA) separated by 90 s inter-tone intervals (ITI). Mice learned to avoid foot-shocks by stepping onto a nearby platform located in the opposite corner of the operant box than the reinforced lever at the cost of losing access to the VI30 scheduled reward. During the extinction phase, mice were exposed to the same contextual and auditory cues in the operant chambers, but foot-shocks were not delivered. Lever presses responses and magazine entries were recorded using MedPC IV software and freezing and time in the platform (avoidance) automatically scored by ANY-maze (Stoelthing Co.).

### Statistics

No statistical power estimation analyses were used to predetermine sample sizes, which instead were chosen to match previous publications ^11,17,60^. All statistics were performed in R v4.2.2 mostly relying on *stats v4.0.2*, *tidyverse v1.3.1* and *lmerTest v3.1-3* packages. In summary, pairwise comparisons were performed with Welch’s *t*-tests (*stats::t.test* function), correlations using Pearson’s *r* (*stats::cor.test* function), independence testing with χ^2^ tests (*stats::chisq.test* function) and more complex multifactorial designs were analyzed using linear models computed with the *stats::lm* function for fixed effects-only models or *lmerTest::lmer* function for mixed effects models. Random effects (conceptualizing non-independent observations, such as repeated measures and/or nested observations) were modeled as random intercept factors. Subsequent analysis of variance (LMM-ANOVA) was performed using type III sums of squares with Kenward-Roger’s approximation of degrees of freedom. Post-hoc testing was performed using the *emmeans* package and significance was adjusted for false discovery rate (FDR) at a 0.05 level using standard Benjamini-Hochberg procedure ^62^. Bar and line graphs represent mean ± sem. Correlation graphs represent regression line with its 95% confidence interval. Significance was set at *p* < 0.05.

### Data availability

All snRNAseq data reported in this study are deposited in the Gene Expression Omnibus under accession number GSE227313. All other data, including raw photometry data, are deposited in a Github repository. *Both will be made public upon publication - for review, please email the corresponding author for private access keys*.

### Code availability

Custom R scripts and code utilized in this study, including for statistical analysis, are available upon request.

#### Acknowledgments

The authors would like to thank Stephen Pirpinias, Katherine Beach, Catherine McManus, Kyra Schmidt, Nathalia Pulido and Ezekiell Mouzon for transgenics breeding and genotyping, and XuQiang Qiao and Dr. Edgardo Aritzia from the Dean’s Flow Cytometry CoRE at the Icahn School of Medicine at Mount Sinai for assistance in nuclei sorting. This work was supported by grants from the Boehringer Ingelheim Fonds (PhD Fellowship to A.G.) and the National Institutes of Health (RF1MH128970 and U01MH116442 to P.R., R01DA014133 and R01MH051399 to E.J.N.)

## Author information

### Contributions

A.G. and E.J.N. conceived the project. A.G. developed all methodology, except for the PMA task that was developed by A.M.M.T and F.J.M.R. A.G. performed all behavioral experiments and photometry recordings, with help from A.M.M.T for PMA testing and E.M.P. for stereotaxic surgeries. M.S. performed nuclei sorting and confocal imaging. V.P. performed electrophysiological recordings, under R.D.B. supervision. J.F.F. performed snRNAseq library preparation. P.R. contributed resources for snRNAseq and software for analysis. C.M. contributed software for photometry analysis. A.G. performed all formal analysis and visualization. A.G. wrote the manuscript, which was reviewed and edited by M.S., A.M.M.T., J.F.F., F.J.M.R., C.M., P.R. R.D.B. and E.J.N.

### Corresponding authors

Correspondence to Eric J. Nestler.

---

**Extended Data Fig. 1:**
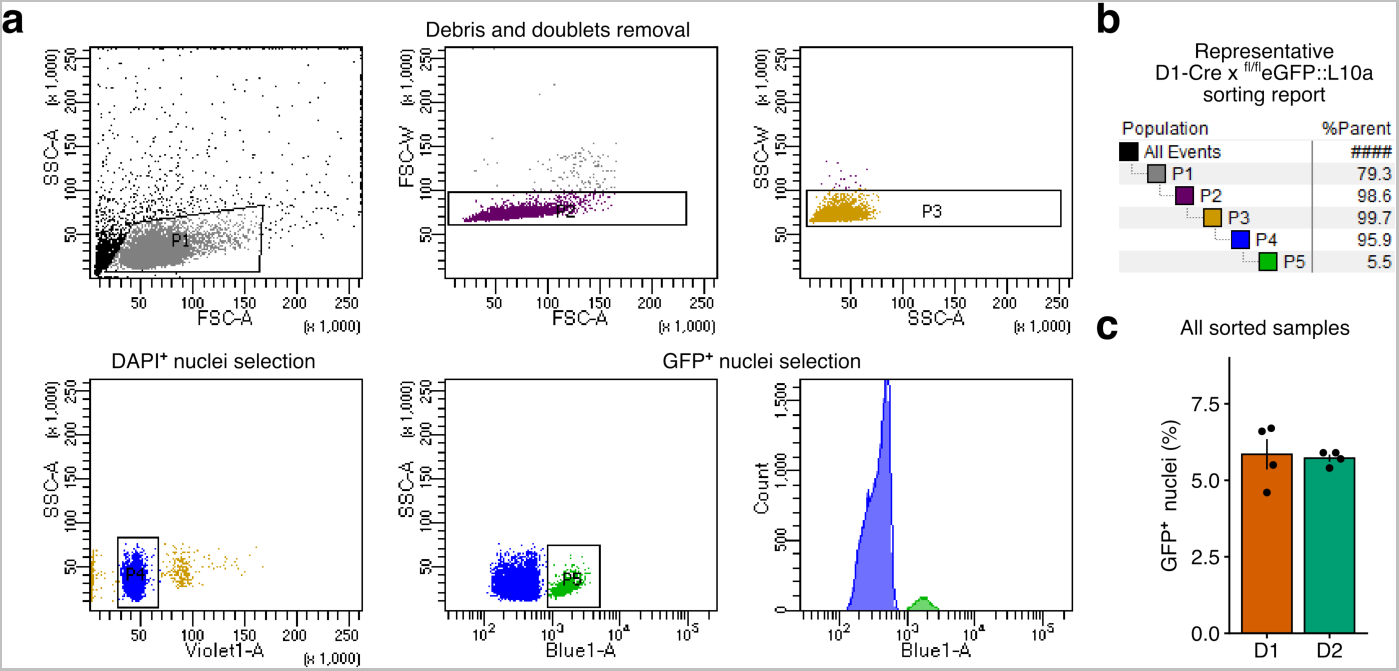
Fluorescence-Activated Nuclei Sorting (FANS) of D1 and D2 vHipp cells. **a**, Representative FANS gating strategy from a D1-Cre x ^fl/fl^eGFP::L10a male sample. **b**, Summary of FAN-sorting hierarchical gating strategy. **c**, Percent of GFP-positive nuclei for all D1-Cre and D2-Cre x ^fl/fl^eGFP::L10a sorted samples.

**Extended Data Fig. 2:**
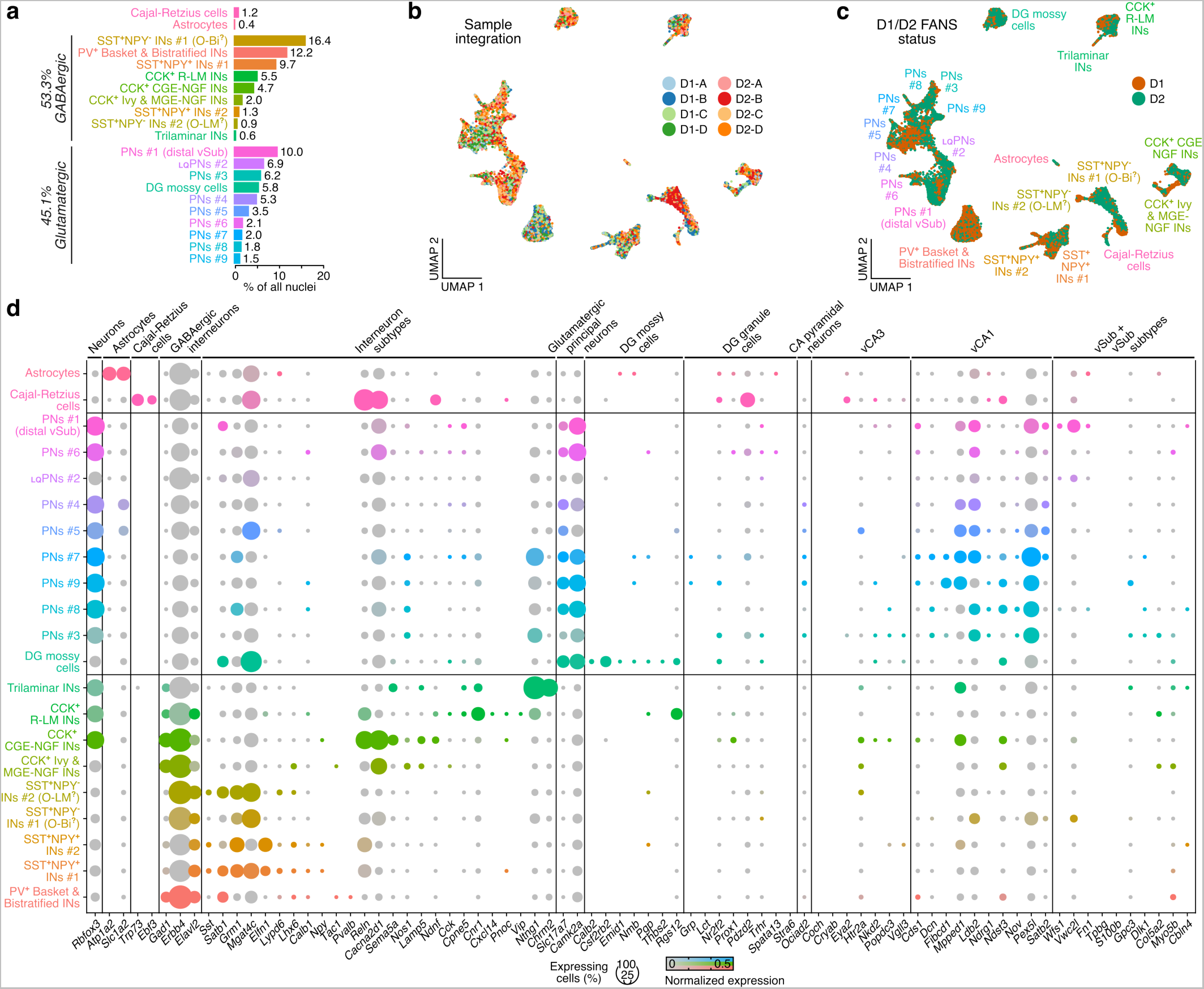
Sample integration and cell-type annotation of snRNAseq clusters. **a**, Proportion of all nuclei in individual clusters. **b**, Sample integration highlighting intermingled contribution of nuclei originating in each sample to clusters. **c**, D1-sorted or D2-sorted origin of individual nuclei, quantified in Fig. 2c. **d,** Expression of published marker genes for different cell types across clusters. Note the absence of cluster enriching for marker genes of DG granule cells and CA3 pyramidal neurons. Full lists of cluster marker genes available in Supplementary Information Table 1.

**Extended Data Fig. 3:**
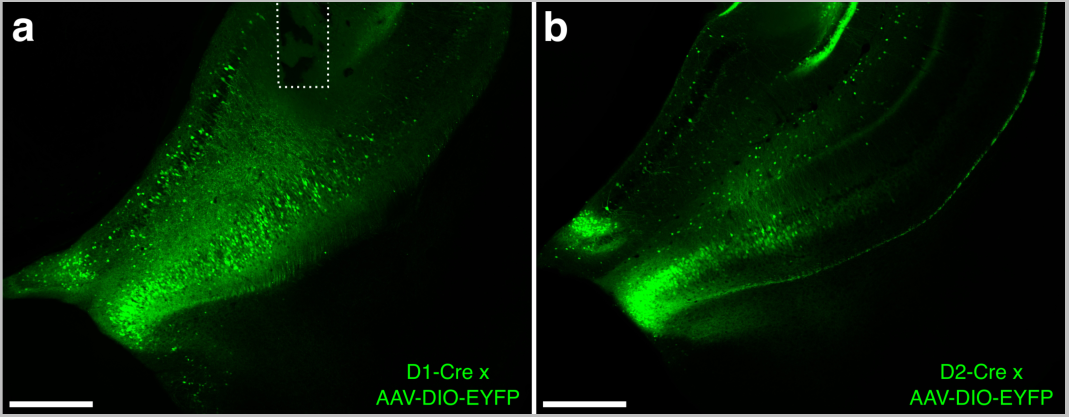
Viral targeting of vSub. **a**, Representative image of AAV-DIO-EYFP expression in the vSub of a D1-Cre male mouse. Dotted line depicts placement of an optogenetic fiber optic. **b**, Representative image of AAV-DIO-EYFP expression in the vSub of a D2-Cre male mouse. Animals with significant somatic expression in neighboring entorhinal cortex (<10%) were removed from analysis. Scale bars 500 µm.

**Extended Data Fig. 4:**
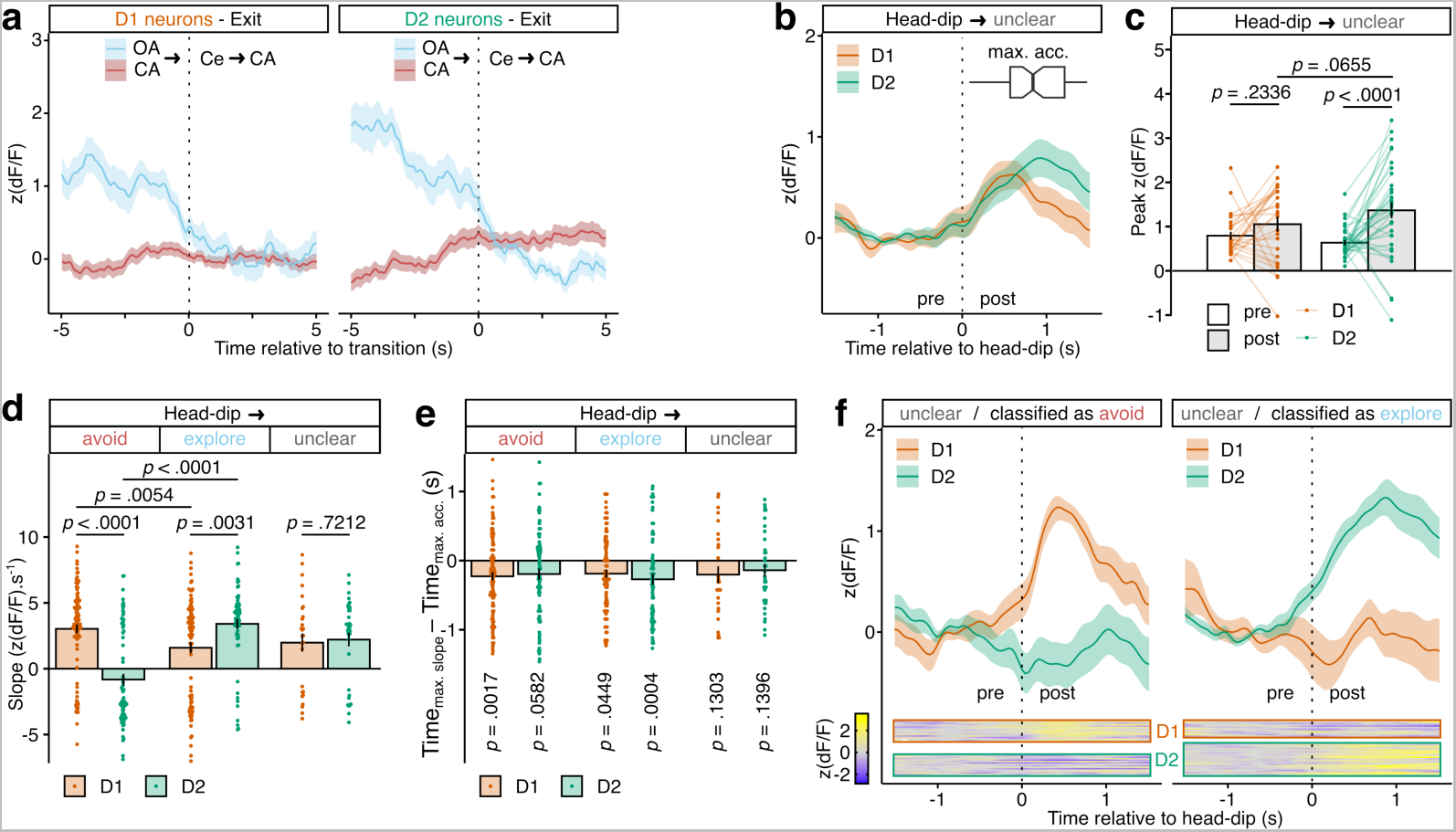
Additional analyses of calcium imaging of vHipp D1 and D2 neuronal activity during EPM testing. **a**, Average GCaMP6s signal in D1-Cre (left) and D2-Cre (right) mice during exits from the open arm (OA, blue) or closed arm as a control (CA, red). Only traces when the mouse did a complete open/closed arm to center (Ce) to closed arm are used for averaging. **b**, Average GCaMP6s signal in D1-Cre (red) and D2-Cre (green) mice around head-dips outside the EPM apparatus event with unclear outcome. Boxplot represents the median time ± inter-quartile range of the times of maximal acceleration (max. acc.) after each head-dip, indicating movement initiation. **c**, Quantification of maximum (peak) GCaMP6s signal before (pre) or after (post) each head-dip event (statistics in Fig. 3). **d**, Quantification of GCaMP6s signal change slope after head-dip events. LMM-ANOVA: cell type F_1,23.29_ = 1.29 *p* = 0.2679, outcome F_2,380.68_ = 6.90 *p* = 0.0011, cell type x outcome F_2,380.68_ = 28.50 *p* < 0.0001; followed by FDR-adjusted post-hoc tests. **e**, Quantification of the time delay between the time of maximal GCaMP6s signal change (max. slope) and the time of movement initiation (max. acc.). LMM-summary: intercept ≠ 0 t_67.07_ = −3.68 *p* = 0.0005, and LMM-ANOVA: cell type F_1,23.10_ = 0.0053 *p* = 0.9427, outcome F_2,383.84_ = 0.19 *p* = 0.8241, cell type x outcome F_2,383.84_ = 0.46 *p* = 0.6310; followed by individual LMM-summary for intercept ≠ 0 for each cell type x outcome combination and FDR adjustment of p-values. **f**, Classification of unclear-outcome head-dips using a SVM trained on manually-annotated avoid/explore D1 and D2 time-series. Both average GCaMP6s signal (top) and heatmaps of individual time-series (bottom) are represented for both D1 and D2 signals, split by SVM outcome prediction. Data represented as mean ± sem.

**Extended Data Fig. 5:**
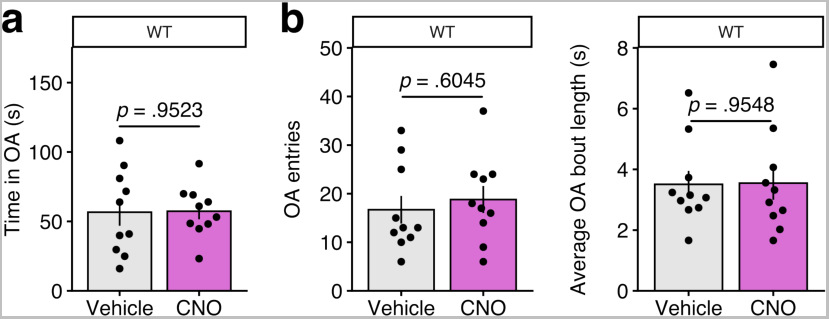
Effects of CNO itself on EPM behavior. Wild-type male mice were injected i.p. with CNO (3 mg/kg, n = 10) or vehicle (n = 10) 15 min before EPM testing. **a**, Quantification of open arm exploration time. Welch’s t-test: t_14.65_ = −0.061 *p* = 0.9523. **b**, Quantification of the total number of open arm entries (left; Welch’s t-test: t_17.99_ = −0.53 *p* = 0.6045) and of the average time of each open arm exploration bout (right; Welch’s t-test: t_17.26_ = −0.057 *p* = 0.9548).

**Extended Data Fig. 6:**
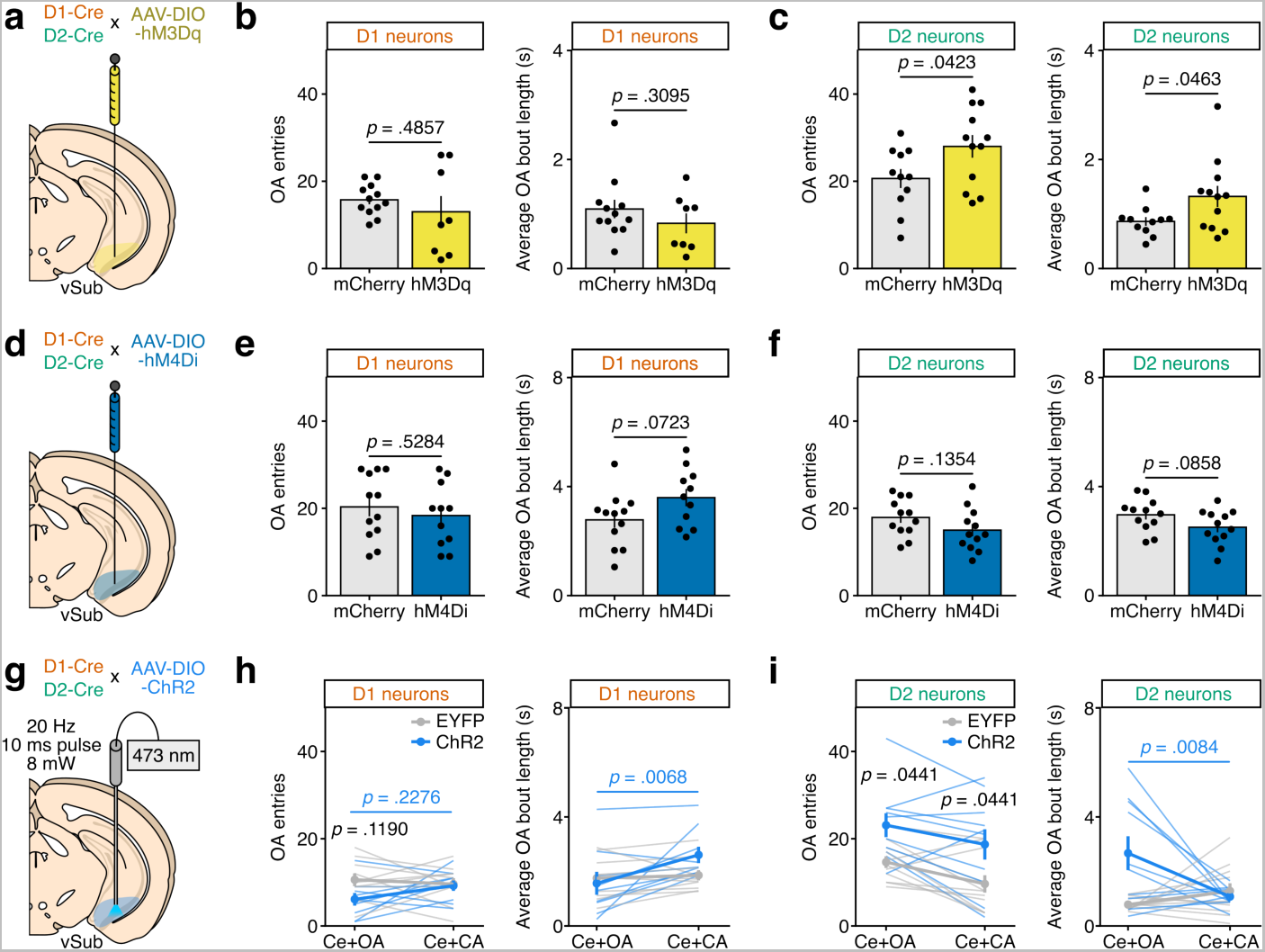
Additional analyses of chemogenetic and optogenetic manipulation of vHipp D1 and D2 neurons during EPM testing. **a**, Experimental schematic. Male D1-Cre and D2-Cre mice were injected in vSub with an AAV-DIO-hM3Dq (n = 8 D1, n = 12 D2) or a control AAV-DIO-mCherry (n = 12 D1, n = 11 D2). CNO (3 mg/kg) was administered i.p. to all animals 15 min before testing. **b**, Quantification of the total number of open arm entries (left; Welch’s t-test: t_8.16_ = 0.73 *p* = 0.4857) and of the average time of each open arm exploration bout (right; Welch’s t-test: t_16.42_ = 1.05 *p* = 0.3095) for D1-Cre mice. **c**, Quantification of the total number of open arm entries (left; Welch’s t-test: t_20.68_ = −2.16 *p* = 0.0423) and of the average time of each open arm exploration bout (right; Welch’s t-test: t_14.63_ = −2.18 *p* = 0.0463) for D2-Cre mice. **d**, Experimental schematic. Male D1-Cre and D2-Cre mice were injected in vSub with an AAV-DIO-hM4Di (n = 11 D1, n = 12 D2) or a control AAV-DIO-mCherry (n = 12 D1, n = 12 D2). CNO (3 mg/kg) was administered i.p. to all animals 15 min before testing. **e**, Quantification of the total number of open arm entries (left; Welch’s t-test: t_20.94_ = 0.64 *p* = 0.5284) and of the average time of each open arm exploration bout (right; Welch’s t-test: t_20.60_ = −1.89 *p* = 0.0723) for D1-Cre mice. **f**, Quantification of the total number of open arm entries (left; Welch’s t-test: t_21.68_ = 1.55 *p* = 0.1354) and of the average time of each open arm exploration bout (right; Welch’s t-test: t_21.93_ = 1.80 *p* = 0.0858) for D2-Cre mice. **g**, Experimental schematic. Male D1-Cre and D2-Cre mice were injected in vSub with an AAV-DIO-ChR2 (n = 9 D1, n = 10 D2) or a control AAV-DIO-EYFP (n = 10 D1, n = 11 D2). Optogenetic stimulation (473 nm laser, 8 mW, 20 Hz, 10 ms pulses) was delivered when the animal was either in the center or open arm (Ce+OA) or in the center and closed arm (Ce+CA) in a within-subject design. **h**, Quantification of the total number of open arm entries (left; LMM-ANOVA: stimulation zone F_1,17_ = 0.67 *p* = 0.4232, virus F_1,17_ = 2.63 *p* = 0.1234, stimulation zone x virus F_1,17_ = 2.55 *p* = 0.1284; followed by FDR-adjusted post-hoc tests) and of the average time of each open arm exploration bout (right; LMM-ANOVA: stimulation zone F_1,17_ = 8.88 *p* = 0.0084, virus F_1,17_ = 0.58 *p* = 0.4580, stimulation zone x virus F_1,17_ = 5.8529 *p* = 0.0271; followed by FDR-adjusted post-hoc tests) for D1-Cre mice. **i**, Quantification of the total number of open arm entries (left; LMM-ANOVA: stimulation zone F_1,19_ = 8.70 *p* = 0.0082, virus F_1,19_ = 7.94 *p* = 0.0110, stimulation zone x virus F_1,19_ = 0.036 *p* = 0.8526; followed by FDR-adjusted post-hoc tests) and of the average time of each open arm exploration bout (right; LMM-ANOVA: stimulation zone F_1,19_ = 2.62 *p* = 0.1221, virus F_1,19_ = 6.1645 *p* = 0.0225, stimulation zone x virus F_1,19_ = 9.55 *p* = 0.0060; followed by FDR-adjusted post-hoc tests) for D2-Cre mice. Data represented as mean ± sem.

**Extended Data Fig. 7:**
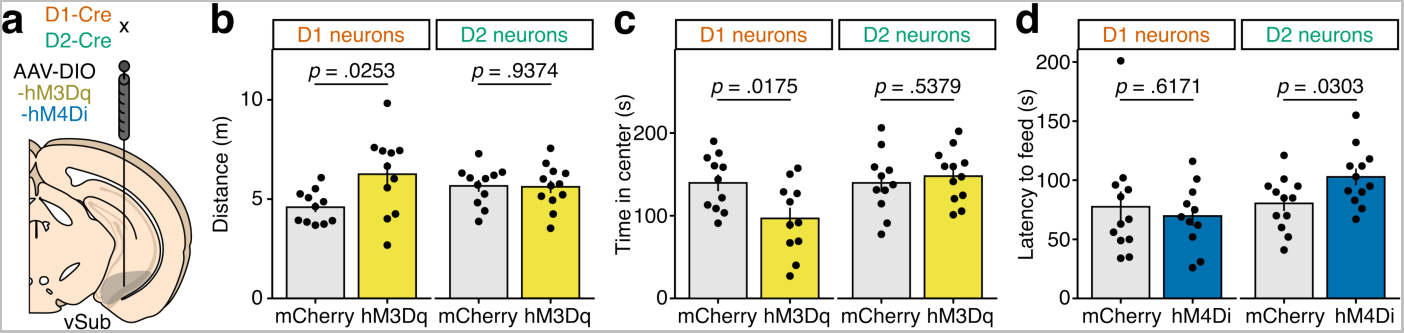
Chemogenetic manipulation of vHipp D1 and D2 neurons during anxiety-related testing. **a**, Experimental schematic. Male D1-Cre and D2-Cre mice were injected in vSub with an AAV-DIO-hM3Dq (n = 11 D1, n = 12 D2) or a control AAV-DIO-mCherry (n = 11 D1, n = 11 D2), and another cohort was injected with either an AAV-DIO-hM4Di (n = 11 D1, n = 12 D2) or a control AAV-DIO-mCherry (n = 11 D1, n = 12 D2). CNO (3 mg/kg) was administered i.p. to all animals 15 min before testing in an open-field test (OFT) for the first cohort and for novelty-suppressed feeding (NSF) for the second. **b**, Quantification of total locomotor activity in OFT for D1-Cre (left; Welch’s t-test: t_13.25_ = −2.52 *p* = 0.0253) and D2-Cre (right; Welch’s t-test: t_20.99_ = 0.08 *p* = 0.9374) mice. **c**, Quantification of time spent in the center of an OFT for D1-Cre (left; Welch’s t-test: t_18.84_ = 2.6052 *p* = 0.0175) and D2-Cre (right; Welch’s t-test: t_19.62_ = −0.63 *p* = 0.5379) mice. **d**, Quantification of the latency to the first feeding bout during NSF testing for D1-Cre (left; Welch’s t-test: t_18.31_ = 0.51 *p* = 0.6171) and D2-Cre (right; Welch’s t-test: t_21.82_ = −2.32 *p* = 0.0303) mice. Data represented as mean ± sem.

**Extended Data Fig. 8:**
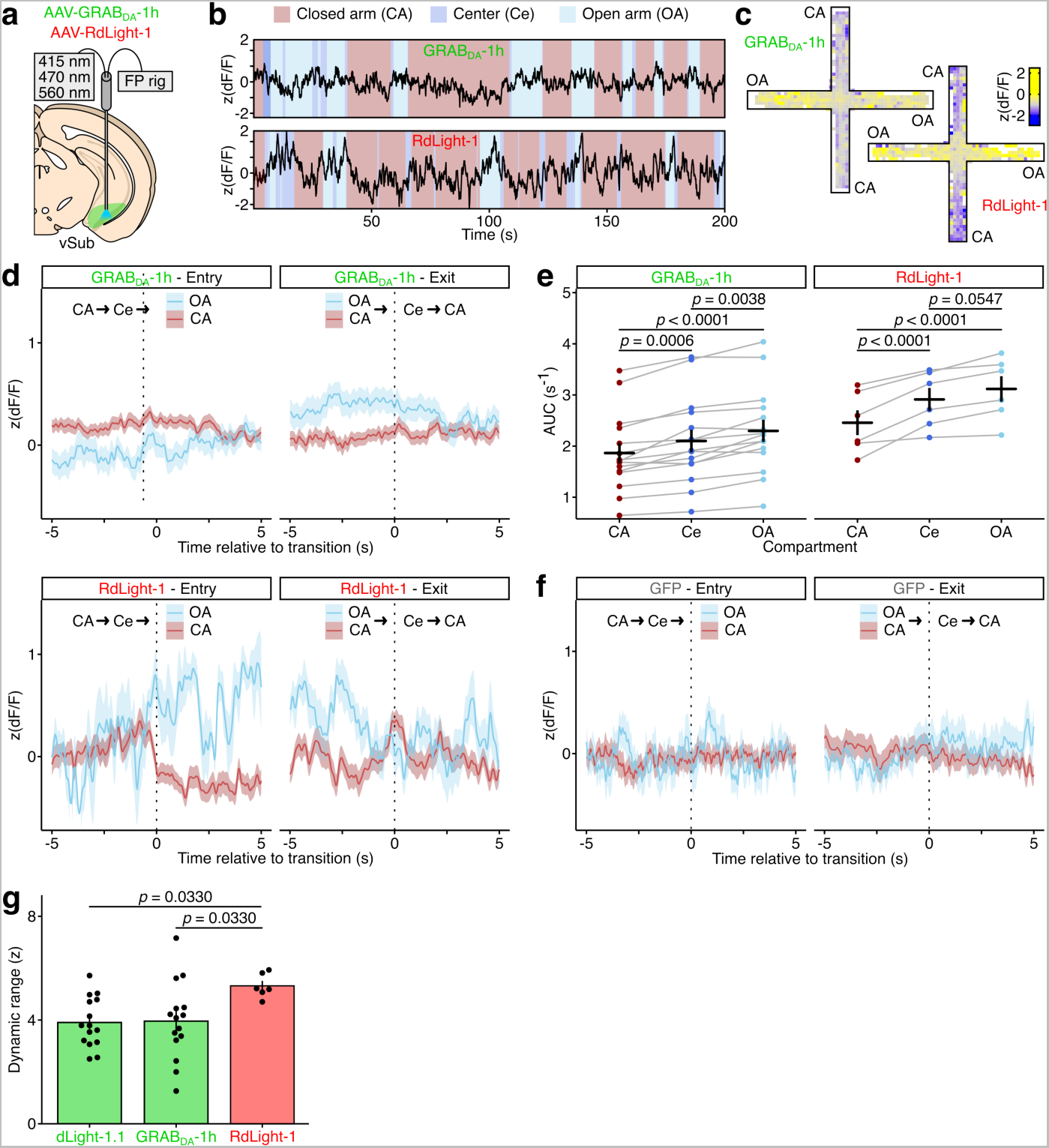
*In vivo* dopamine sensing in vSub – other sensors and sensor comparisons. **a**, Experimental schematic. Male mice were injected in vSub with an AAV-dLight-1.1 (n = 15, Fig. 5), an AAV-GRAB_DA_-1h (n = 15), an AAV-RdLight-1 (n = 6) or with a control AAV-GFP (n = 8, Fig. 5) and implanted with optical fibers before recording during EPM testing. **b**, Representative traces during EPM exploration for a GRAB_DA_-1h (top) and RdLight-1 (bottom) animal. **c**, Spatially averaged fluorescence signal in the EPM for all GRAB_DA_-1h (top-left) and all RdLight-1 (bottom-right) mice. **d**, Average GRAB_DA_-1h (top) and RdLight-1 (bottom) signal during entries to (left) and exits from (right) the open arm (OA, blue) or closed arm as a control (CA, red). Only traces when the mouse did a complete closed arm to center (Ce) to open/closed arm are used for averaging. **e**, Average area under the curve (AUC) by EPM compartment for GRAB_DA_-1h (left) and RdLight-1 (right) mice. LMM-ANOVA: compartment F_2,80_ = 49.82 *p* < 0.0001, sensor F_3,40_ = 1.70 *p* = 0.1822, compartment x sensor F_6,80_ = 5.1529 *p* = 0.0002; followed by FDR-adjusted post-hoc tests. **f**, Average control GFP signal during entries to (left) and exits from (right) the OA (blue) or CA as a control (red). **g**, Quantification of *in vivo* dynamic range for all three dopamine sensors used in this study, calculated as the difference between maximum and minimum z(dF/F) reached during recording for each animal. LMM-ANOVA: sensor F_2_ = 3.51 *p* = 0.0414; followed by FDR-adjusted post-hoc tests. Data represented as mean ± sem.

**Extended Data Fig. 9:**
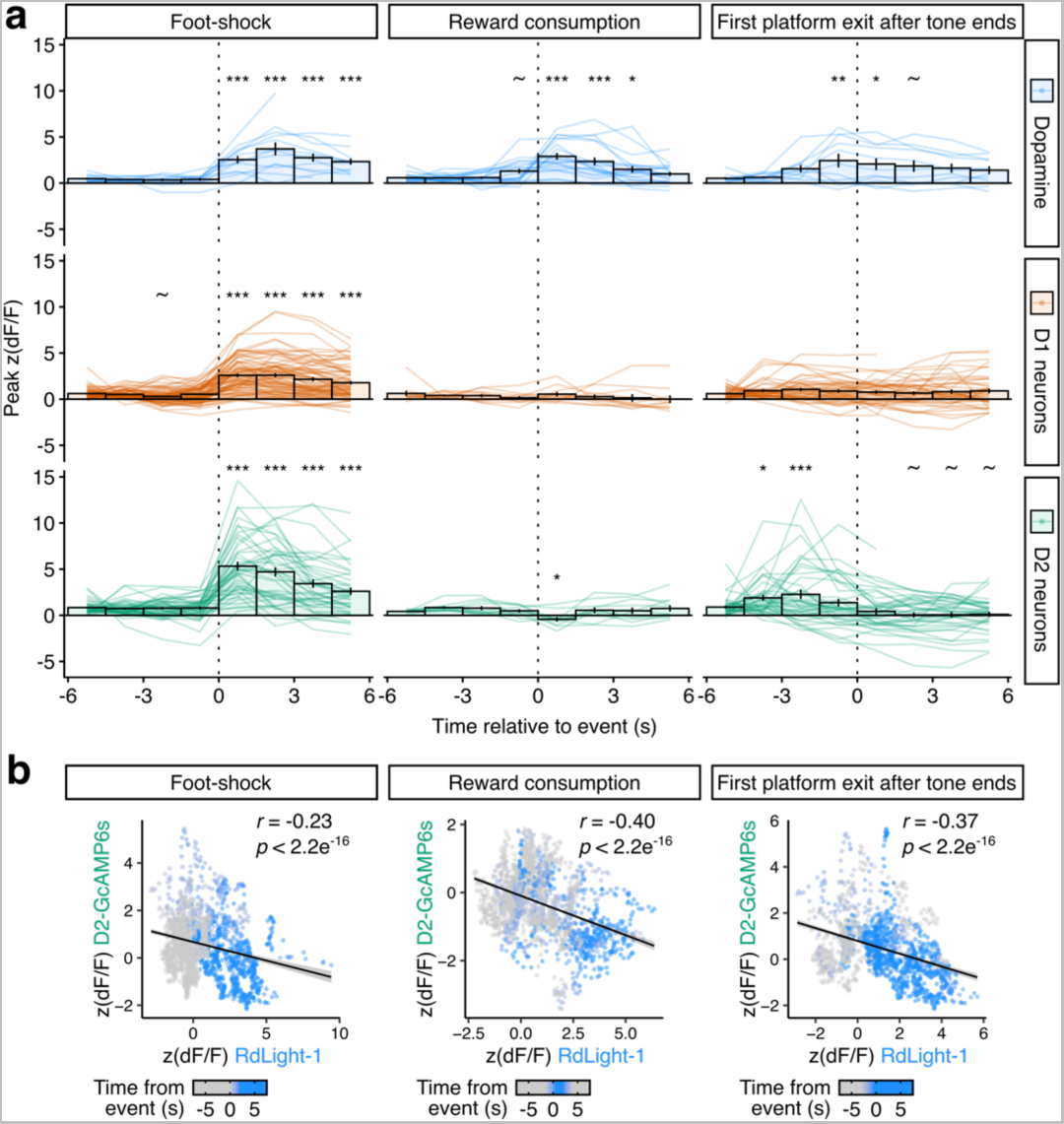
vSub dopamine, D1 and D2 correlates of approach and avoidance in the PMA task. **a**, Quantification of peri-event photometry signal changes shown in Fig. 6, measured as the maximum (peak) z(dF/F) value reached within successive 1.5 s-long time-bins. All were analyzed with LMM-ANOVA for the time-bin factor and followed by FDR-adjusted post-hoc tests. All pairwise comparisons were computed but only the ones significant *versus* the first 1.5 s time-bin are represented for clarity (~ p < 0.1, * p < 0.05, ** p < 0.01, *** p < 0.001). Foot-shock, DA: F_7,80.86_ = 19.69 *p* < 0.0001. Foot-shock, D1: F_7,548.52_ = 66.88 *p* < 0.0001. Foot-shock, D2: F_7,302.07_ = 46.81 *p* < 0.0001. Reward, DA: F_7,133_ = 12.29 *p* < 0.0001. Reward, D1: F_7,70_ = 0.67 *p* = 0.7005. Reward, D2: F_7,70_ = 3.09 *p* = 0.0068. First platform exit, DA: F_7,70_ = 3.57 *p* = 0.0024. First platform exit, D1: F_7,311.15_ = 0.91 *p* = 0.4989. First platform exit, D2: F_7,262.51_ = 9.9281 *p* < 0.0001. **b**, Correlation between RdLight-1 and D2-GCaMP6s signals in one D2-Cre mouse injected in vSub with both an AAV-DIO-GCaMP6s and an AAV-RdLight-1 and recorded with dual color photometry, around foot-shocks (left; Pearson’s *r* = −0.23 *p* < 0.0001), reward consumption (middle; Pearson’s *r* = −0.40 p < 0.0001) and first platform exit after tone end (right; Pearson’s *r* = −0.37, *p* < 0.0001). For bar graphs, data represented as mean ± sem, with shaded lines representing individual events. For correlation graphs, regression lines are shown with their 95% confidence intervals.

**Extended Data Fig. 10:**
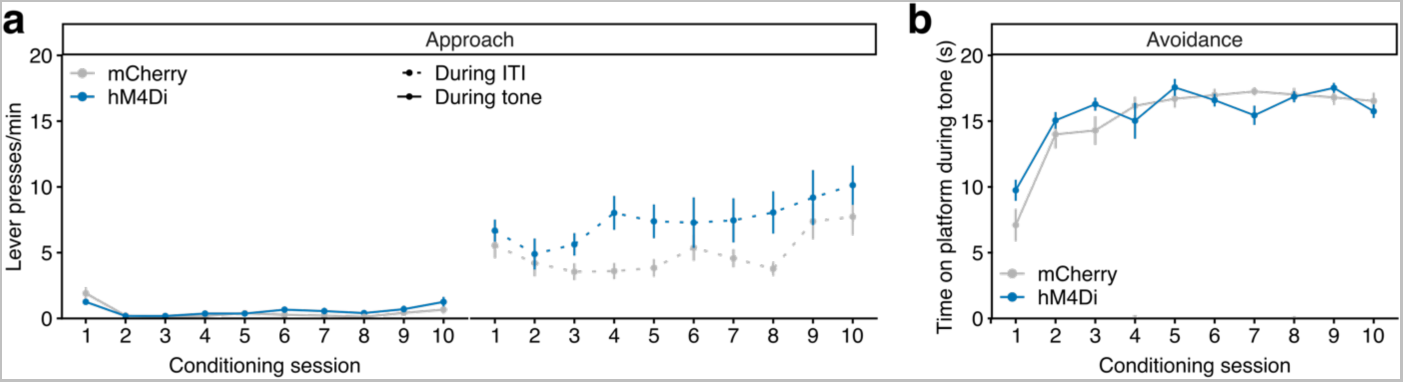
Chemogenetic manipulation of vSub D2 neurons in the PMA task – acquisition behaviors. Male D2-Cre mice were injected in vSub with an AAV-DIO-hM4Di (n = 9) or a control AAV-DIO-mCherry (n = 11). **a**, Approach behavior during conditioning sessions, measured as the lever pressing rate in responses per minute during tone presentation (left; LMM-ANOVA: session F_9,162_ = 10.30 *p* < 0.0001, DREADD F_1,18_ = 1.03 *p* = 0.3227, session x DREADD F_9,162_ = 1.54 *p* = 0.1373) or during inter-tone intervals (ITI; right; LMM-ANOVA: session F_9,162_ = 5.50 *p* < 0.0001, DREADD F_1,18_ = 4.12 *p* = 0.0575, session x DREADD F_9,162_ = 1.11 *p* = 0.3589). **b**, Avoidance behavior during conditioning sessions, measured as the time spent on the platform during the 20 s tone. LMM-ANOVA: session F_9,162_ = 26.31 *p* < 0.0001, DREADD F_1,18_ = 0.44 *p* = 0.5170, session x DREADD F_9,162_ = 1.91 *p* = 0.05385. Data represented as mean ± sem.

